# State-selective Modulation of Heterotrimeric Gαs Signaling with Macrocyclic Peptides

**DOI:** 10.1101/2020.04.25.054080

**Authors:** Shizhong A. Dai, Qi Hu, Rong Gao, Hayden Peacock, Emily E. Blythe, Ziyang Zhang, Mark von Zastrow, Hiroaki Suga, Kevan M. Shokat

**Affiliations:** Department of Cellular and Molecular Pharmacology, University of California San Francisco, San Francisco, CA, 94158, USA; Howard Hughes Medical Institute; Department of Chemistry, Graduate School of Science, The University of Tokyo, 7-3-1 Hongo, Bunkyo-ku, Tokyo 113-0033, Japan; Department of Psychiatry, University of California, San Francisco, San Francisco, CA, 94158, USA

## Abstract

The G protein-coupled receptor (GPCR) cascade leading to production of the second messenger cAMP is replete with pharmacologically targetable receptors and enzymes with the exception of the Gα subunit, Gαs. GTPases remain largely undruggable given the difficulty of displacing high-affinity guanine nucleotides and the lack of other drug binding sites. We explored a chemical library of 10^12^ cyclic peptides in order to expand the chemical search for inhibitors of this enzyme class. We identified two macrocyclic peptides, GN13 and GD20, that antagonize the active and inactive states of Gαs, respectively. Both macrocyclic peptides fine-tune Gαs activity with high nucleotide-binding-state selectivity and G protein class-specificity. Co-crystal structures reveal that GN13 and GD20 distinguish the conformational differences within the switch II/α3 pocket and block effector interactions. The Gαs inhibitors showed strong activity in cellular contexts through binding to crystallographically defined pockets. The discovery of cyclic peptide inhibitors targeting Gαs provides a path for further development of state-dependent GTPase inhibitors.

## INTRODUCTION

The family of human GTPases represents a vast but largely untapped source of pharmacological targets. They serve as key molecular switches that control cell growth and proliferation through cycling between tightly regulated ON/OFF states. The role of specific GTPase family members across diverse human diseases have been widely established by cancer genome sequencing (e.g., *KRAS, GNAS* and others) and by familial studies in neurodegenerative disease (e.g. *LRRK2, RAB39B*) (Prior et al., 2012; O’Hayre et al., 2013; Alessi and Sammler, 2018; Wilson et al., 2014). Despite the widespread recognition of these disease target relationships, only very recently has the first drug targeting a GTPase K-Ras (G12C) achieved clinical proof of principle (Canon et al., 2019; Hallin et al., 2020). The covalent somatic cysteine mutant-specific nature of the K-Ras (G12C) drugs has opened the potential for targeting a GTPase for the first time.

Several peptide-based probes that non-covalently target GTPases have been reported, but they either lack proper drug-like properties or have limited target scope (Takasaki, et al., 2004; Ja and Roberts, 2004; Johnston et al., 2005; Johnston et al., 2005; Johnston et al., 2006; Ja et al., 2006; Austin et al., 2008). Short linear peptides have shown good ability to target the switch II/α3 helix region in the heterotrimeric G protein α-subunit (Gα) with high nucleotide binding state selectivity. However, linear peptides are not the ideal molecules for drug discovery because of their poor cell-permeability and inherent instability in cells.

Cyclic peptides are promising candidates for GTPase drug development. Like linear peptides, cyclic peptides are also capable of targeting protein-protein interfaces (Sohrabi et al., 2020). Peptide cyclization stabilizes the peptide sequence and constrains the flexible peptide conformations for better cell penetration (Dougherty et al., 2019). Cyclic peptide inhibitors of Gα proteins have been reported: for instance, the macrocyclic depsipeptide natural product YM-254890 targets GDP-bound Gαq with high specificity and potency in cells (Nishimura et al., 2010). Despite the highly conserved structure of G proteins and the recent chemical tractability of fully synthetic YM-254890, efforts to use this natural macrocycle as a scaffold from which to discover inhibitors of other G proteins (Gαs, Gαi) have not been successful (Kaur et al., 2015; Xiong et al., 2016; Zhang et al., 2017), likely because of the limited chemical diversity of available YM-254890 analogs. We therefore reasoned that screening an ultra-large chemical library of cyclic peptides against a given nucleotide binding state of Gαs might allow us to discover Gαs nucleotide-binding-state-selective inhibitors that discriminate between the active and inactive states of Gαs and potentially open the remainder of the GTPase family to pharmacological studies.

The Random nonstandard Peptide Integrated Discovery (RaPID) system (Yamagishi et al., 2011) is an *in vitro* display system which merges the flexibility of an *in vitro* translation system (FIT) (Murakami et al., 2006; Murakami et al., 2003., Ramaswamy et al., 2004; Xiao et al., 2008) with mRNA display, enabling the screening of exceptionally large macrocyclic peptide libraries (> 10^12^ molecules) against challenging targets (Passioura and Suga, 2017). Here we report the discovery by the RaPID system of GN13 and GD20, two macrocyclic peptides that are the first known cell-permeable, nucleotide-state-selective inhibitors of Gαs, with high selectivity over other G protein subfamilies and nucleotide binding states.

## RESULTS

### Selection of state-selective cyclic peptides that bind to the active or inactive state of Gαs

Affinity screening hits emerging from the RaPID cyclic peptide selection process against Gαs could theoretically bind anywhere on the surface of the protein and so might or might not perturb its function. To increase the probability of selecting a function-perturbing hit, we took advantage of the fact that when Gαs switches from the GDP-bound inactive state to the GTP-bound active state significant conformational changes occur on one face of Gαs, comprising the so-called switch I, II and III regions (Lambright et al., 1994), which are known to bind inhibitor or effector protein partners such as Gβγ or adenylyl cyclases (Tesmer et al., 1997; Liu et al., 2019) (Figure 1A). We reasoned that performing both a positive selection against one state of Gαs and a negative selection against the other state would enrich for binders to the switch regions, and that these binders would be likely to state-selectively disrupt Gαs function.

**Figure 1.**
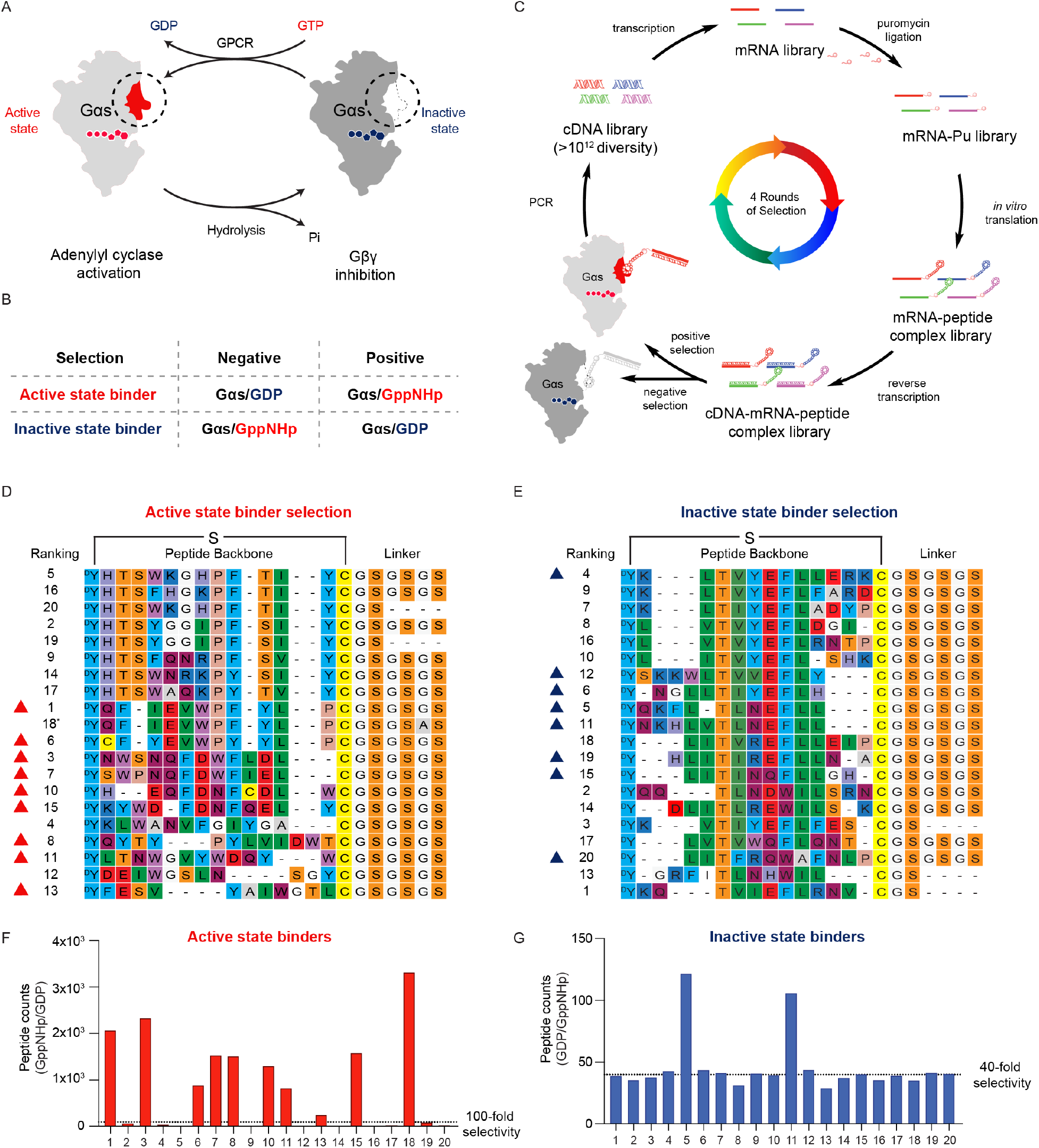
RaPID selection of state-selective Gαs binding cyclic peptides. (A) The molecular switch Gαs adopts distinct conformations, governed by its guanine nucleotide binding state. Switch regions are highlighted with circle. (B) A selection strategy to achieve state-selectivity of Gαs binders. (C) Schematic representation of RaPID selection. (e.g., Gαs active state binder selection, positive selection = Gαs/GppNHp (light grey), negative selection = Gαs/GDP (dark grey)). The mRNA library was ligated with puromycin, translated, and reverse-transcribed to yield our peptide-mRNA-cDNA complex library, which was subjected to sequential negative and positive selections. Negative selection was not included in the first round of selection. DNA sequences of cyclic peptide binders were identified by PCR. (D-E) Sequence alignment of top 20 cyclic peptides from the last round of positive selections. The sulfur bridge cyclizes peptides between D-tyrosine and cysteine. The 18th peptide (asterisk) from the active state binder selection was not selected because it has the same sequence as the 1st peptide except a Gly to Ala mutation in the linker region. (F-G) Comparison selection was performed by analyzing peptide-mRNA-cDNA complex binding to GDP-bound or GppNHp-bound Gαs-immobilized beads from the last round of selections, respectively. A high ratio means a better selectivity between the positive/negative selection. Cyclic peptides with high selectivity are marked with triangles in (D) or (E) and were selected for solid phase synthesis. See also Figure S1.

In order to select a Gαs active state binding peptide, we performed the positive selection with wild-type Gαs (WT Gαs) bound to the non-hydrolyzable GTP analogue GppNHp (5′-guanylyl imidodiphosphate) and the negative selection against GDP-bound WT Gαs. A parallel Gαs inactive state binder selection was performed using GDP-bound WT Gαs as the positive selection and GppNHp-bound WT Gαs as the negative selection (Figure 1B). Starting from a cDNA library, each round of selection included PCR amplification of the cDNA library, *in vitro* transcription into an mRNA library, ligation with a puromycin linker, and translation to generate a peptide library covalently conjugated with its encoding mRNA library (Figure 1C). The library encoded peptides contain an N-chloroacetyl-D-tyrosine at the N-terminus, followed by 8-12 random proteinogenic amino acids encoded by NNK codons, a cysteine residue and a GSGSGS linker (Figure 1D and 1E). Cyclization occurs spontaneously between the chloroacetyl group and the thiol group of the downstream cysteine residue (or possibly those appeared in the random region). The peptide-ligated mRNA library was further reverse-transcribed into a cDNA-mRNA-peptide library, subjected to a negative selection against one state of Gαs, then followed by a positive selection against the other state of Gαs (Figure 1C).

After four rounds of selection (R1-R4), cyclic peptide binders for the GppNHp-bound or GDP-bound Gαs were enriched (Figure S1A and S1B) and identified by next generation sequencing (NGS). The sequences of the top 20 hits were aligned and shown in Figure 1D and 1E. Selective cyclic peptides from the R4 pool were identified and characterized by comparison selection against the respective positive and negative protein baits (Figure 1F and 1G, see also Figure S1C). Nine of the top twenty hits from the active state binder selection (with more than 100-fold selectivity for GppNHp-bound over GDP-bound Gαs, indicated by red triangles in Figure 1D) and eight of the top twenty hits from the inactive state binder selection (with more than 40-fold selectivity for GDP-bound over GppNHp-bound Gαs, indicated by blue triangles in Figure 1E) were chosen for further analysis. To evaluate the cyclic peptide hits without the appended DNA/mRNA duplex, residues from the N-chloroacetyl-D-tyrosine to the glycine (after the anchor cysteine residue) of the selected peptides were synthesized using solid phase peptide synthesis followed by *in situ* cyclization.

### Active state binding cyclic peptide GN13 blocks Gαs-mediated adenylyl cyclase activation

In order to determine whether (Gαs/GppNHp) specific binders inhibit Gαs activity, we assayed the ability of Gαs to activate its downstream effector adenylyl cyclase (AC) (Figure 2A). We refer to resynthesized active state binding cyclic peptides with a “GN” (Gαs/GppNHp) preceding their ranking number. We first tested the physical interaction between GppNHp-bound Gαs and adenylyl cyclase in the presence of the active sate binders using a fluorescence resonance energy transfer (FRET) assay (Figure S2A). Eight out of nine GN peptides showed strong, dose-dependent inhibition of the interaction between Gαs and adenylyl cyclase (Figure 2B). We then evaluated the inhibitory effect of the top hits on Gαs-mediated adenylyl cyclase activation by measuring production of cAMP in a reconstituted Gαs activity assay (Figure 2A). Among these nine active state binders, GN13 showed the greatest inhibition, with an IC_50_ of 4.15 ± 1.13 μM (Figure 2C and 2D). GN13 did not inhibit forskolin-mediated, Gαs-independent adenylyl cyclase activation (Figure S2B), suggesting a Gαs-dependent mechanism of inhibition.

**Figure 2.**
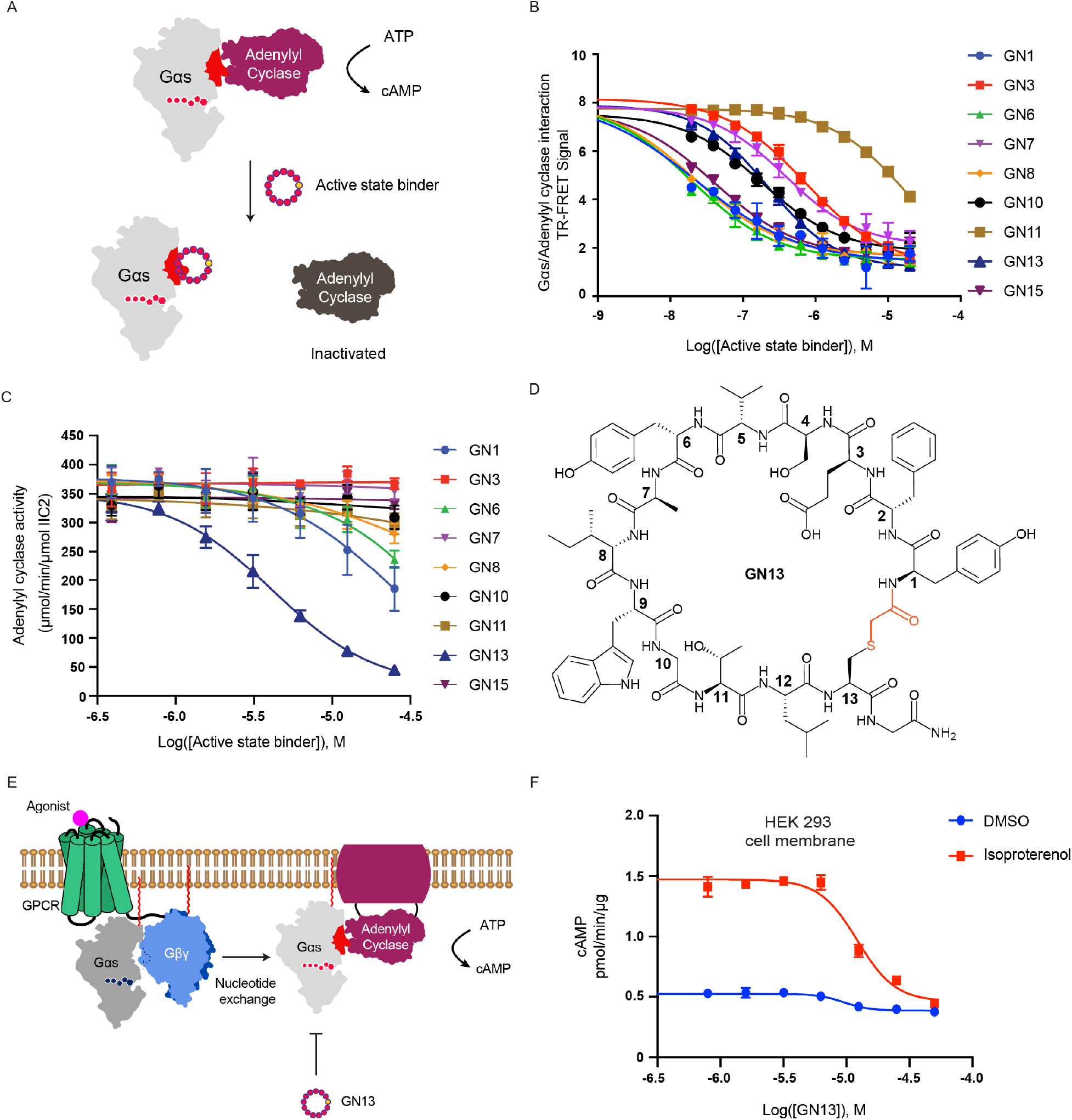
Gαs active state inhibitor GN13 inhibits Gαs-mediated adenylyl cyclase activation. (A) Schematic representation of active state binders inhibiting Gαs mediated adenylyl cyclase activation. (B) Activation of adenylyl cyclase by Gαs was inhibited by active state binders in a dose-dependent manner. GppNHp-bound Gαs was mixed with various concentrations of cyclic peptides, adenylyl cyclase (VC1/IIC2), and forskolin. After adding ATP, the reaction was carried out at 30 °C for 10 min. Production of cAMP was evaluated by the LANCE Ultra cAMP kit. The data represent the mean ± SE of three independent replicates. (C) Active state binders inhibited the protein-protein interaction between biotinylated Gαs WT and His-tagged adenylyl cyclase in a dose-dependent manner. The data represent the mean ± SD of three independent replicates. (D) Chemical structure of the resynthesized cyclic peptide GN13. Cyclization linkage was highlighted (red). (E) Schematic representation of GN13 inhibiting GPCR-stimulated Gαs activity in cell membrane. (F) Cell membranes were prepared from HEK293 cells and were preincubated with GTP/GDP mixture (500/50μM) and various concentrations of GN13 for 2hours, and then stimulated with 40 µM of β2AR agonist isoproterenol. After adding ATP, the reaction was carried out at 30 °C for 30 min. Production of cAMP was evaluated by the LANCE Ultra cAMP kit. The data represent the mean ± SD of three independent replicates. See also Figure S2.

The inhibitory Gα protein, Gαi, is a negative regulator in the cAMP pathway and possesses a structure similar to that of Gαs (Gilman., 1987). To assess whether GN13 was capable of discriminating between Gαs and Gαi, we measured the binding kinetics of GN13 to Gαi and Gαs using biolayer interferometry (BLI). We found that GN13 binds to immobilized GppNHp-bound Gαs with a *K*_D_ value of 0.19 ± 0.02 μM (Figure S2C). By contrast, GN13 showed no detectable binding to GDP-bound Gαs, GDP-bound Gαi1 or GppNHp-bound Gαi1 at the highest concentration tested (Figure S2D, S2E, and S2F), confirming the state-selectivity and class-specificity of GN13 for the active state of Gαs.

We sought to test the efficacy of GN13 in β2-adrenergic receptor (β2AR) mediated second messenger stimulation in cell membranes. Membrane anchored GDP-bound Gαs is inhibited by Gβγ in the resting state. It can undergo GPCR mediated GDP to GTP exchange upon agonist stimulation (Weis and Kobilka, 2018). The presence of GN13 could potentially capture the newly generated GTP-bound Gαs and inhibit its activation (Figure 2E). To test this idea, we incubated cell membranes prepared from HEK293 cells with GN13 and examined cAMP accumulation with or without stimulation of β2-adrenergic receptor (β2AR) by isoproterenol (ISO). GN13 inhibited ISO-stimulated Gαs activity to a background level, with an IC_50_ of 12.21 ± 2.51 μM (Figure 2F). These results suggested that GN13 can modulate Gαs activation by agonist stimulated β2AR.

### The crystal structure of GppNHp-bound Gαs in complex with GN13

To elucidate how the cyclic peptide GN13 binds to Gαs and inhibits Gαs-mediated adenylyl cyclase activation, we solved a co-crystal structure of the Gαs/GppNHp/GN13 complex. The structure was determined by molecular replacement and refined to 1.57 Å (Table S1). The overall structure is shown in Figure 3A. GN13 assumes a highly ordered structure through extensive intramolecular and intermolecular hydrogen bonding networks with three well-defined water molecules (Figure S3A-C). One molecule of GN13 binds to a pocket between switch II and the α3 helix of GppNHp-bound Gαs through hydrogen bonding and hydrophobic interactions (Figure 3B and 3C). Specifically, the side chain of residue E3 in GN13 accepts a hydrogen bond from the ε-amino group of K274 in Gαs; the indole ring of residue W9 in GN13 donates a hydrogen bond to the side chain of E268 in Gαs; the main chain carbonyl oxygens of residues V5, W9 and T11, and the main chain amide of W9 in GN13 form five hydrogen bonds with residues N279, R280, R231, R232, and S275 on switch II and the α3 helix in Gαs (Figure 3B). The side chains of residues I8 and W9 (IW motif) in GN13 dock into two hydrophobic pockets on Gαs (Figure 3C), giving rise to a high Gαs binding affinity.

**Figure 3.**
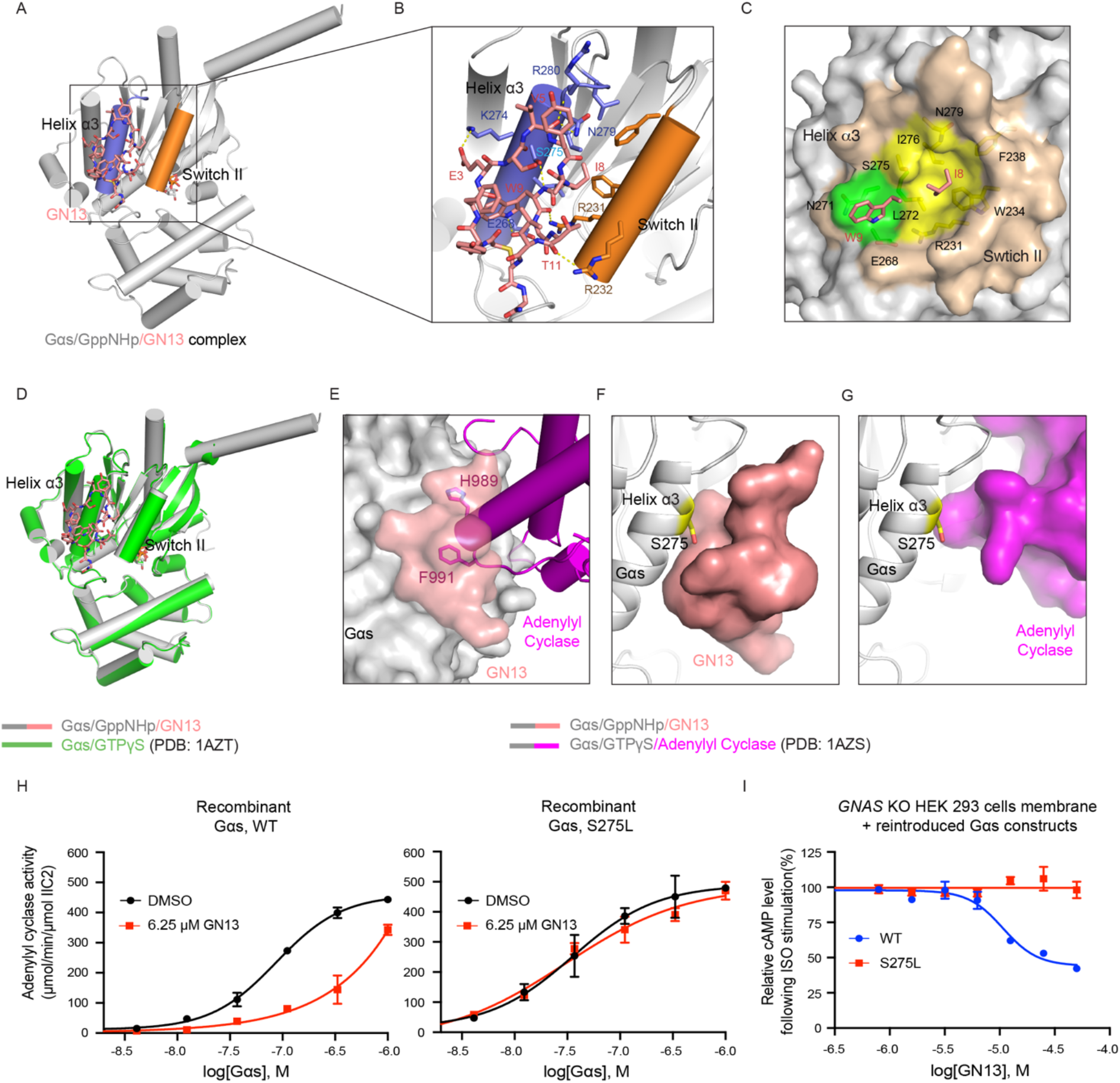
Crystal Structure of GppNHp-bound Gαs in complex with GN13. (A) Overall structure of the GN13/GppNHp-bound Gαs complex. GN13 (salmon) binds in between the switch II region (orange) and the α3 helix (slate). GN13 and GppNHp are shown as sticks. (B) Structural details of the GN13-Gαs interaction. Ion pair and hydrogen bonds are represented by yellow dashed lines. (C) Close-up view of two hydrophobic pockets (green and yellow) that accommodate the tryptophan and isoleucine side chains of GN13 (salmon). Residues that form those pockets are depicted as stick models and labeled. (D) Alignment of Gαs/GN13 complex structure (grey) with the structure of GTPγS-bound Gαs (green) (PDB: 1AZT). Root mean square deviation (RMSD) = 0.479 Å. (E) Gαs/GN13 (grey/salmon) complex structure was aligned with the structure of GTPγS-bound Gαs/adenylyl cyclase complex (grey/magenta) (PDB: 1AZS). GN13 blocks H989/F991 of adenylyl cyclase from binding to the same pocket in Gαs. (F-G) Close-up view of the interaction between GN13 (salmon) and the Gαs α3 helix (grey) in (F) and the interaction between adenylyl cyclase (magenta) and the Gαs α3 helix in (G) (PDB: 1AZS). S275 is shown as sticks. (H) WT Gαs and the S275L mutant have comparable biochemical activities in the adenylyl cyclase activation assay in the presence of Gβ1/γ2 (black curves). 6.25 μM of GN13 inhibits adenylyl cyclase activation by Gαs WT (red curve, left) but not by Gαs S275L (red curve, right). The data represent the mean ± SD of three independent measurements. (I) The Gαs S275L mutation confers resistance to GN13 inhibition in HEK293 cell membranes. *GNAS* KO HEK 293 cells were transiently transfected with Gαs WT (blue) or S275L mutant (red) constructs and followed by cell membrane preparation. Cell membranes were treated with various concentrations of GN13 for 2hours, and then stimulated with 40 µM of β2AR agonist isoproterenol. After adding ATP, the reaction was carried out at 30 °C for 30 min. Production of cAMP was evaluated by the LANCE Ultra cAMP kit. The data represent the mean ± SD of three independent replicates. See also Figure S3.

Structural analysis reveals that residue W9 in GN13 is centrally located within the interface between GN13 and Gαs, contributing three hydrogen bonds as well as hydrophobic interactions with the switch II/α3 pocket (Figure 3B and 3C). Analogous to this key tryptophan in GN13, PDEγ (effector of G protein transducin, Gt) residue W70 (Slep et al., 2001) and adenylyl cyclase II (effector of Gαs) residue F991 (Tesmer et al., 1997) are inserted into the same switch II/α3 clefts of Gt and Gαs (Figure 3E, see also Figure S3E). Comparison between the Gαs/GppNHp/GN13 complex structure and a Gαs/GTPγS/adenylyl cyclase complex structure (PDB: 1AZS) suggested that GN13 directly occludes the hydrophobic interaction between Gαs and adenylyl cyclase (Tesmer et al., 1997), which accounts for the inhibitory effect of GN13 (Figure 3E).

### Structural basis for the nucleotide state-selectivity and the G protein class-specificity of GN13

The Gαs/GppNHp/GN13 complex strongly resembles the Gαs/GTPγS structure (PDB: 1AZT) (Sunahara et al., 1997), suggesting that GN13 recognizes the active state conformation and does not induce significant conformational change upon binding (Figure 3D). We aligned our structure with the structure of inactive GDP-bound Gαs (chain I in PDB 6EG8) (Liu et al., 2019). In comparison with our structure, the N-terminus of switch II in the GDP-bound Gαs is unstructured and close to the α3 helix, with nearly half of the GN13/Gαs interface disrupted (Figure S3F). In particular, R232 of switch II (shown in space filling) is predicted to create a steric clash with I8 of GN13. Therefore, the GN13-bound complex structure explains the state-selectivity of GN13 for the active state of Gαs.

GN13 showed excellent G protein class-specificity, although we did not include other Gα proteins, such as Gαi, as a part of the negative selection. To identify the G-protein specificity determinants of GN13, we aligned our structure with the structure of active state Gαi in complex with a Gαi-specific linear peptide, KB1753 (PDB: 2G83) (Johnston et al., 2006). The affinity determinant, IW motif, was presented in both GN13 and KB1573, but Gαs-specific binding of GN13 was mainly determined by charge interactions (Figure 3B, see also Figure S3G and S3H): (1) D229 in Gαs (homologous residue in Gαi: S206) confines the orientation of switch II R232, which forms a hydrogen bond with a main chain carbonyl oxygen of T11 in GN13. (2) K274 in Gαs (homologous residue in Gαi: D251) interacts with the negatively charged side chain of E3 in GN13. Homologous residues in Gαi either do not engage with or even repulse against GN13, rendering the lack of GN13 binding.

To further assess the cellular specificity of GN13, we designed a GN13-resistant Gαs mutant based on structural analysis. We examined the structures of the Gαs/GN13 complex and Gαs/AC complex and noticed that serine 275 in Gαs makes close contact with GN13, but does not contact adenylyl cyclase (Figure 3F and 3G). We reasoned that mutating this serine residue to a bulky residue should create a drug-resistant Gαs mutant that blocks the GN13-Gαs interaction but would have little effect on adenylyl cyclase activation. Indeed, Gαs S275L mutant maintained a similar level of biochemical activity but was resistant to GN13 inhibition (Figure 3H, see also Figure S3I). We tested GN13 in the membranes of *GNAS*-knockout (*GNAS*-KO) HEK 293 cells that did not express endogenous Gαs protein (Stallaert et al., 2017). GN13 was able to inhibit isoproterenol-mediated cAMP production in cell membranes in a dose-dependent manner when WT Gαs was reintroduced into the *GNAS* KO cells by transient transfection. By contrast, when the drug resistant mutant Gαs S275L was reintroduced into *GNAS* KO cells, the inhibitory effect of GN13 was abolished (Figure 3I), providing further evidence that the observed pharmacological activity is due to GN13 binding to the switch II/α3 pocket in Gαs.

### Inactive state binding cyclic peptide GD20 is a Gαs specific guanine nucleotide dissociation inhibitor (GDI)

The activation of G protein signaling is often rapid and temporary. Gα GTPase activity promotes GTP hydrolysis to GDP and rearranges the switch regions to adopt a GDP-bound inactive conformation. This precisely orchestrated nucleotide binding conformation prevents GDP release, which makes GDP dissociation the rate-limiting step of G protein activation (Dror et al., 2015). An inactive state binder could hypothetically modulate GDP-bound Gαs either by inhibiting GDP release as a guanine nucleotide dissociation inhibitor (GDI) or by facilitating GDP to GTP switch as a guanine nucleotide exchange factor (GEF) (Ghosh et al., 2017). In order to understand how inactive state binders control GDP-bound Gαs function, we carried out a multiple turnover assay to evaluate Gαs steady-state GTPase activity in the presence of top hits from the inactive state binder selection (Figure 4A). The inactive state selected cyclic peptides are indicated with a “GD” (Gαs/GDP) preceding their ranking number, and all of the tested GD peptides showed strong, dose-dependent inhibition against Gαs steady-state GTPase activity. GD20 showed the greatest inhibition, with an IC_50_ of 1.15 ± 0.16 μM (Figure 4B and 4C, see also Figure S4A).

**Figure 4.**
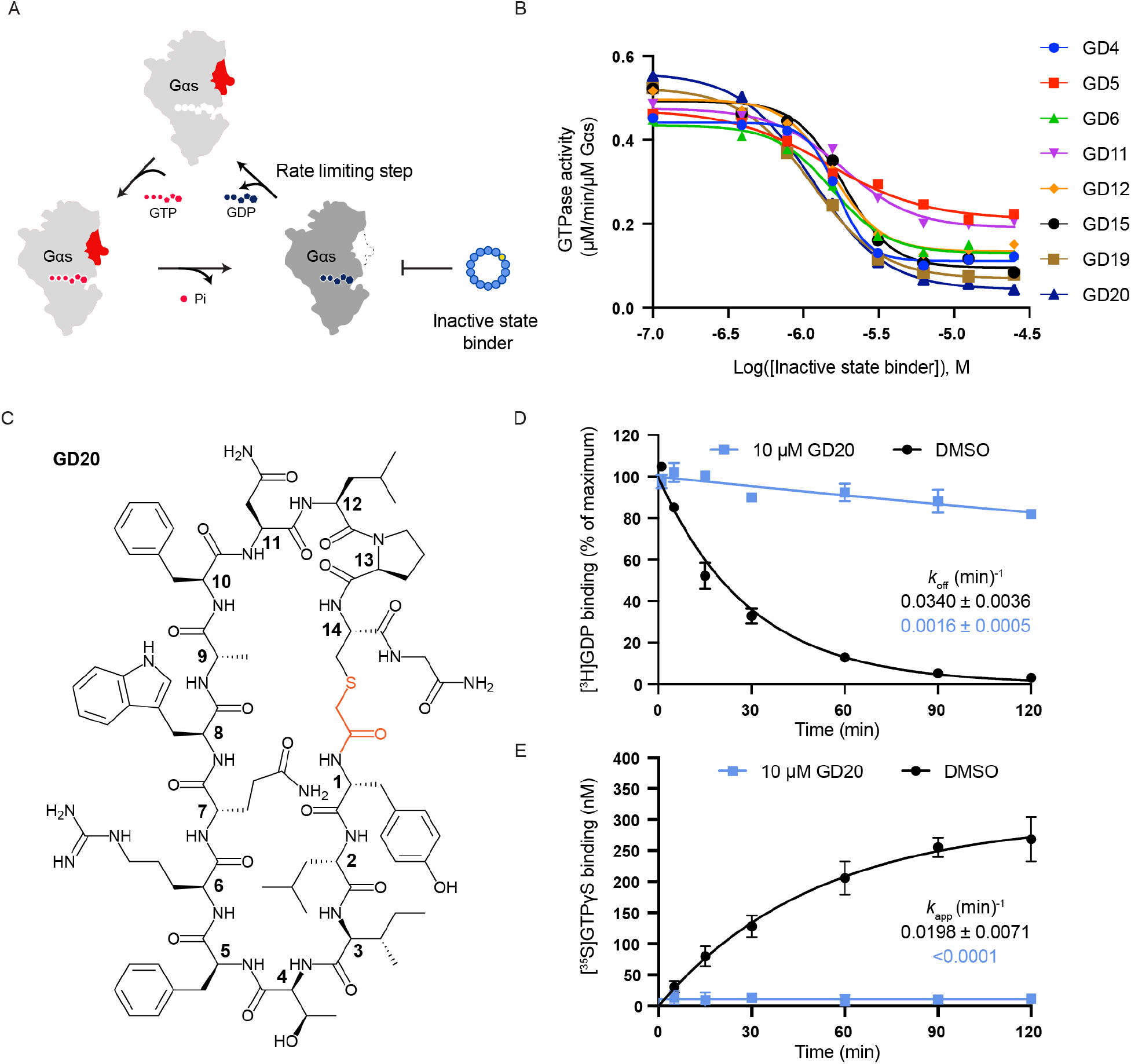
Gαs inactive state inhibitor GD20 inhibits Gαs steady-state GTPase activity by preventing GDP dissociation. (A) Schematic representation of inactive state binders inhibiting Gαs steady-state GTPase activity. (B) Gαs steady-state GTPase activity was inhibited by inactive state binders in a dose-dependent manner. The data represent one measurement. (C) Chemical structure of the resynthesized cyclic peptide GD20. Cyclization linkage was highlighted (red). (D) GD20 slows down the rates of GDP dissociation from Gαs. Gαs preloaded with [^3^H]GDP was assayed in a buffer containing 1 mM MgCl_2_, 0.5 mM GDP, and the indicated concentration of GD20. The data represent the mean ± SD of three independent replicates. (E) The rates of GTPγS binding to Gαs in the presence (blue) or absence (black) of 10 µM GD20 were determined by mixing GDP-bound Gαs with a mixture of [^35^S] GTPγS and GTPγS in a buffer containing 1 mM MgCl_2_. The data represent the mean ± SD of three independent replicates. See also Figure S4.

We determined rate constants for two individual steps in the GTPases cycle, GD20 displayed profound GDI activity towards Gαs by drastically reducing Gαs GDP dissociation rates (*k*_off_) and the apparent rate of GTPγS binding (*k*_app_) to Gαs (Figure 4D and 4E). On the contrary, GN13 only slightly influenced Gαs GDP dissociation, however, increased the apparent rate of GTPγS binding (*k*_app_) to Gαs (Figure S4B and S4C). The discrepancy between GD20, a synthetic Gαs GDI, and GN13, a synthetic Gαs GEF, exemplified how state-selective Gαs binders fine-tune Gαs enzymatic activity. The precise regulation of Gαs by cyclic peptides not only appears at the nucleotide binding state level, but also at the G protein family level. The nucleotide exchange step of a homologous G protein Gαi, is much less sensitive towards GD20 and GN13, confirming the class-specificity of both GD20 and GN13 for Gαs (Figure S4D and S4E).

### The crystal structure of GDP-bound Gαs in complex with GD20

To understand the underlying molecular determinants of how cyclic peptide GD20 favors GDP-bound Gαs and inhibits GDP dissociation. We solved a co-crystal structure of the Gαs/GDP/GD20 complex. The structure was determined by molecular replacement and refined to 1.95 Å (Table S2). The overall structure is shown in Figure 5A. Four well-defined water molecules and a number of intramolecular hydrogen bonds constructed a unique helical secondary structure in GD20 (Figure S5A-E). One molecule of GD20 binds to the cleft between switch II and the α3 helix of GDP-bound Gαs through electrostatic interactions, hydrogen bonding, and hydrophobic interactions (Figure 5B and 5C). Specifically, the side chain of residue R6 in GD20 forms a salt bridge with Gαs E268, and this ion pair is further stabilized by Gαs N271; the main chain carbonyl oxygen of F10 in GD20 forms a hydrogen bond network with S275 and N279 from the α3 helix in Gαs; R231 and W234 on Gαs switch II coordinate a complex hydrogen bond network with W8, N11, L12, C13 and D-tyrosine in GN13; deep inside of the GD20 binding pocket, the main chains of I3 and F5 form multiple hydrogen bonds with G225, Q227, and D229 in Gαs (Figure 5B). These charge and hydrogen bonding interactions rearrange the flexible Gαs switch II motif and bury residues F5 and W8 of GD20 inside of a hydrophobic pocket (Figure 5C). These solidified hydrophobic interactions between GD20 and Gαs likely contribute to the high Gαs binding affinity.

**Figure 5.**
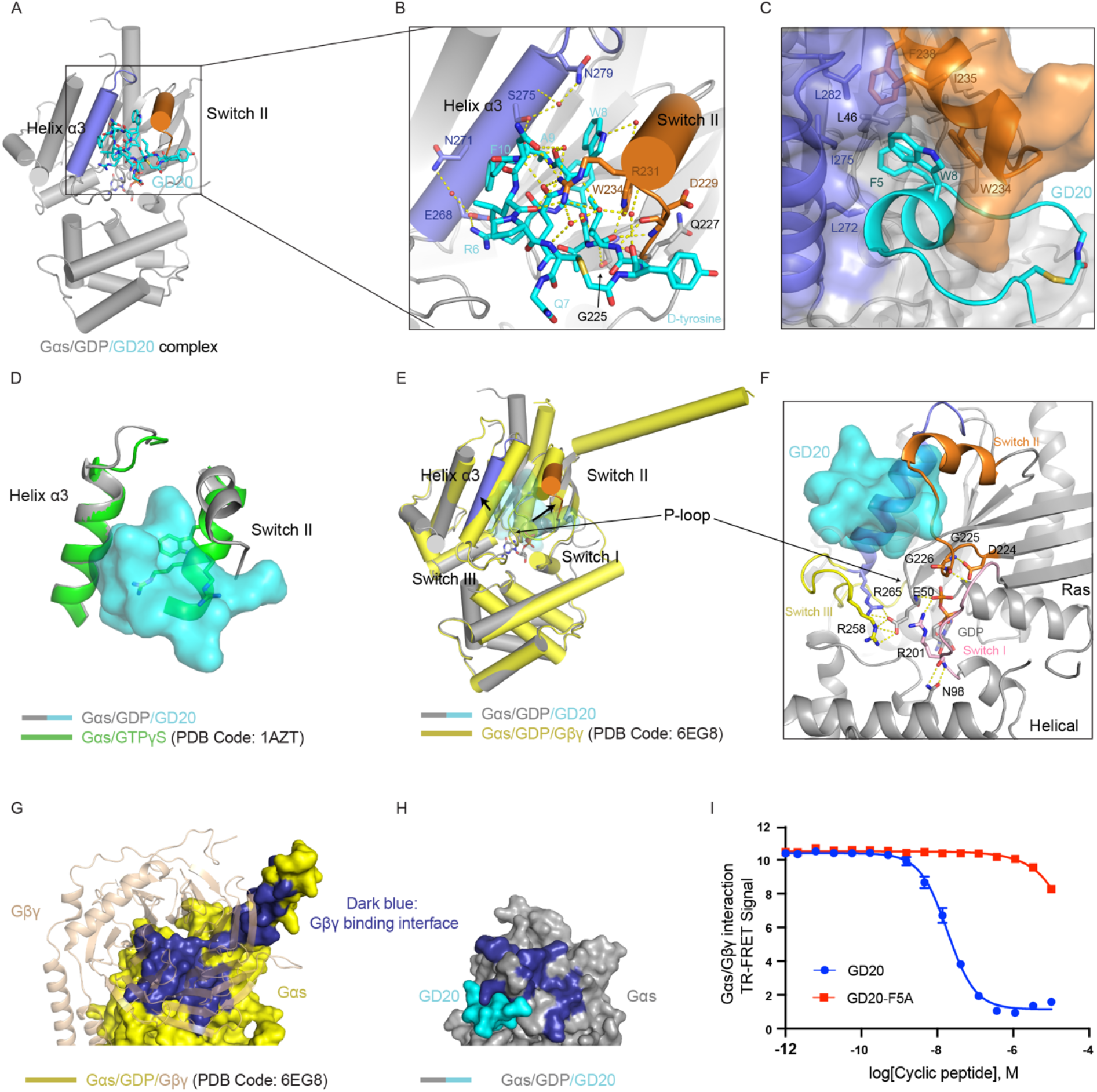
Crystal Structure of GDP-bound Gαs in complex with GD20. (A) Overall structure of the GD20/GDP-bound Gαs complex. GD20 (cyan) binds in between the switch II region (orange) and the α3 helix (slate). GD20 and GDP are shown as sticks. (B) Structural details of the GD20-Gαs interaction. Ion pair and hydrogen bonds are represented by yellow dashed lines. (C) Close-up view of a hydrophobic pocket in Gαs that accommodates the Phe5 and Trp8 side chains of GD20 (cyan). GD20 is shown as cartoon, and Gαs is shown as surface. Residues that form the hydrophobic pocket are depicted as stick models and labeled. (D) Alignment of Gαs/GD20 complex structure (grey) with the structure of GTPγS-bound Gαs (green) (PDB: 1AZT). (E)Alignment of Gαs/GD20 complex structure (grey) with the structure of GDP-bound Gαs (yellow) in the crystal structure of Gαs/Gβ1/γ2 heterotrimer (PDB: 6EG8). Gβγ was hidden for clarity. (F) Close-up view of Gαs nucleotide binding pocket in the Gαs/GD20 complex structure. GD20 is shown as surface. Gαs is shown as cartoon. GDP is shown as sticks. Residues that stabilize GDP binding are depicted as stick models and labeled. Hydrogen bonds are represented by yellow dashed lines. (G) Structural details of Gαs (yellow) and Gβγ (wheat) binding interface (PDB: 6EG8). Gαs is shown as surface, and Gβγ are shown as cartoon. (H) The Gβγ binding interface (dark blue) of Gαs is significantly rearranged when GD20(cyan) binds to Gαs(grey). (I) GD20, but not the GD20-F5A mutant, inhibited the protein-protein interaction between biotinylated Gαs WT and His-tagged Gβγ(C68S) in a dose-dependent manner. The data represent the mean ± SD of three independent replicates. See also Figure S5.

We measured the binding kinetics of GD20 to Gαs using BLI to quantify the binding event. We found that GD20 bound to immobilized GDP-bound Gαs with a *K*_D_ value of 31.4 ± 0.7 nM (Figure S5F). By contrast, GD20 had no detectable binding to GppNHp-bound Gαs, GDP-bound Gαi1 or GppNHp-bound Gαi1 at the highest concentration tested (Figure S5G), confirming the state-selectivity and class-specificity of GD20 for the inactive state of Gαs. A single point mutation F5A nearly completely abolished Gαs binding affinity of GD20, confirming the importance of hydrophobic interactions mediated by F5 (Figure S5H).

### Structural basis for the binding selectivity and biochemical activity of GD20

The high resolution GD20-bound Gαs complex structure elucidated the molecular mechanism by which GD20 distinguishes GDP-bound Gαs over other protein or nucleotide binding states. First, we aligned our GD20-bound Gαs structure with the structure of active GTPγS-bound Gαs (PDB: 1AZT) (Sunahara et al., 1997). The presence of a rigidified switch II motif in the GTPγS-bound Gαs clashes with GD20 (Figure 5D). In particular, R231, R232 and W234 of switch II (shown in space filling) are predicted to create a steric clash with GD20. Second, we compared our GD20-bound Gαs structure with the inactive GDP-bound Gαs structure in complex with Gβγ (chain I in PDB 6EG8) (Figure 5E) (Liu et al., 2019). GD20 binding expands the switch II/α3 pocket by repositioning both motifs. However, structural motifs (such as switch I, III, and the P loop) that are critical for GDP binding remain unchanged with no discernible difference, indicating the GDP-state selective of GD20. Finally, we compared our GD20-bound Gαs structure with the GDP-bound Gαi structure in complex with Gβγ (PDB: 1GP2) (Figure S5I and S3H) (Wall et al., 1995). The specificity of GD20 is determined by three elements which involves electrostatic interactions, Van der Waals interactions, and hydrogen bonding. (1) E268 in Gαs (homologous residue in Gαi: E245) interacts with the positively charged side chain of R6 in GD20. At the same location in Gαi, the presence of a nearby K248 neutralizes the surface charge of Gαi and disfavors the salt bridge formation between Gαi and GD20. (2) F5 and W8 in GD20 dock into a hydrophobic pocket made of two non-polar residues F238 and L282 from Gαs. A L282 to F259 substitution in Gαi structure sterically reshapes the positioning between F238 and L282 in the hydrophobic pocket, dislocates the correct orientation for GD20 Van der Waals interactions. The subtle difference in this pocket between Gαs and Gαi also controls the specificity of other Gα effectors (Chen et al., 2005; Tesmer et al., 2005). (3) The reshaped hydrophobic pockets also influence the residues on the switch II motif. The main chain amide of D229 in Gαs, and the homologous residue S206 in Gαi form different hydrogen bonding patterns. Aspartate 229 in Gαs, but not S206 in Gαi, forms two hydrogen bonds with the main chain amide of I3 in GD20. Homologous residues in Gαi either do not enhance or may even diminish the binding with GD20, resulting in a strong selectivity for Gαs over Gαi.

The GD20-bound Gαs complex structure also demonstrates the molecular basis of GD20 GDI activity (Figure 5F). GDP dissociation from Gαs nucleotide binding site requires both conformational changes that weaken the guanine nucleotide affinity and Ras/Helical domain separation that allows GDP release (Dror et al., 2015). GD20 does not engage the GDP exit tunnel, therefore is not directly occluding GDP release. Instead, GD20 phenocopies the effect of Gβγ GDI activity, rigidifying the juxtapositions of switch I, III, and the P loop. Such a conformational lock not only orients Gαs R201 and E50 to directly capture the β-phosphate of GDP, but also inhibits the spontaneously Ras/Helical domain separation by stabilizing two hydrogen bonds between Gαs R201 and N98. As a result, GD20 strongly antagonizes GDP dissociation from Gαs.

When we compared our GD20-bound Gαs structure with another GDP-bound Gαs structure in complex with Gβγ (Figure 5E) (Liu et al., 2019), we noticed that GD20 induces a significant conformational shift of the Gβγ binding surface, and GD20 occupies the Gβγ binding surface in a competitive manner (Figure 5G and 5H, see also Video S1). We measured the interaction of GDP-bound Gαs and Gβγ in the presence of GD20 or the Gαs binding deficient analog GD20-F5A using a fluorescence resonance energy transfer (FRET) assay (Figure S5J). GD20, but not GD20-F5A, showed a potent, dose-dependent inhibition of the interaction between Gαs and Gβγ, with an IC_50_ of 18.4 ± 2.0 nM (Figure 5I). This inhibitory effect was also Gαs specific, given that GD20 was at least 100-fold more selective for Gαs than Gαi in the FRET assay (Figure S5K). These data suggested that GD20 selectively captures the GDP-bound Gαs, contributing to the Gαs GDI activity and the inhibitory effect against Gαs/Gβγ complex.

### A cell permeable GD20 analog, GD20-F10L, inhibits Gαs/Gβγ interaction in HEK293 cells

Receptor coupled G protein signaling releases GTP-bound Gα and free Gβγ to engage their own effectors to transduce downstream signaling. GDP-bound Gα is a functional “OFF” switch by tightly reassociating with obligate Gβγ dimers and masking the effector binding surfaces on both Gαs and Gβγ (Gulati et al., 2018). A potent Gαs Gβγ protein-protein interaction (PPI) inhibitor should potentially block Gαs Gβγ reassociation and further extend the lifetime of free Gβγ (Figure 6A). With a potent Gαs Gβγ PPI inhibitor, GD20, functioning *in vitro*, we next asked whether GD20 could reduce association of Gαs and Gβγ following receptor stimulation in the cells.

**Figure 6.**
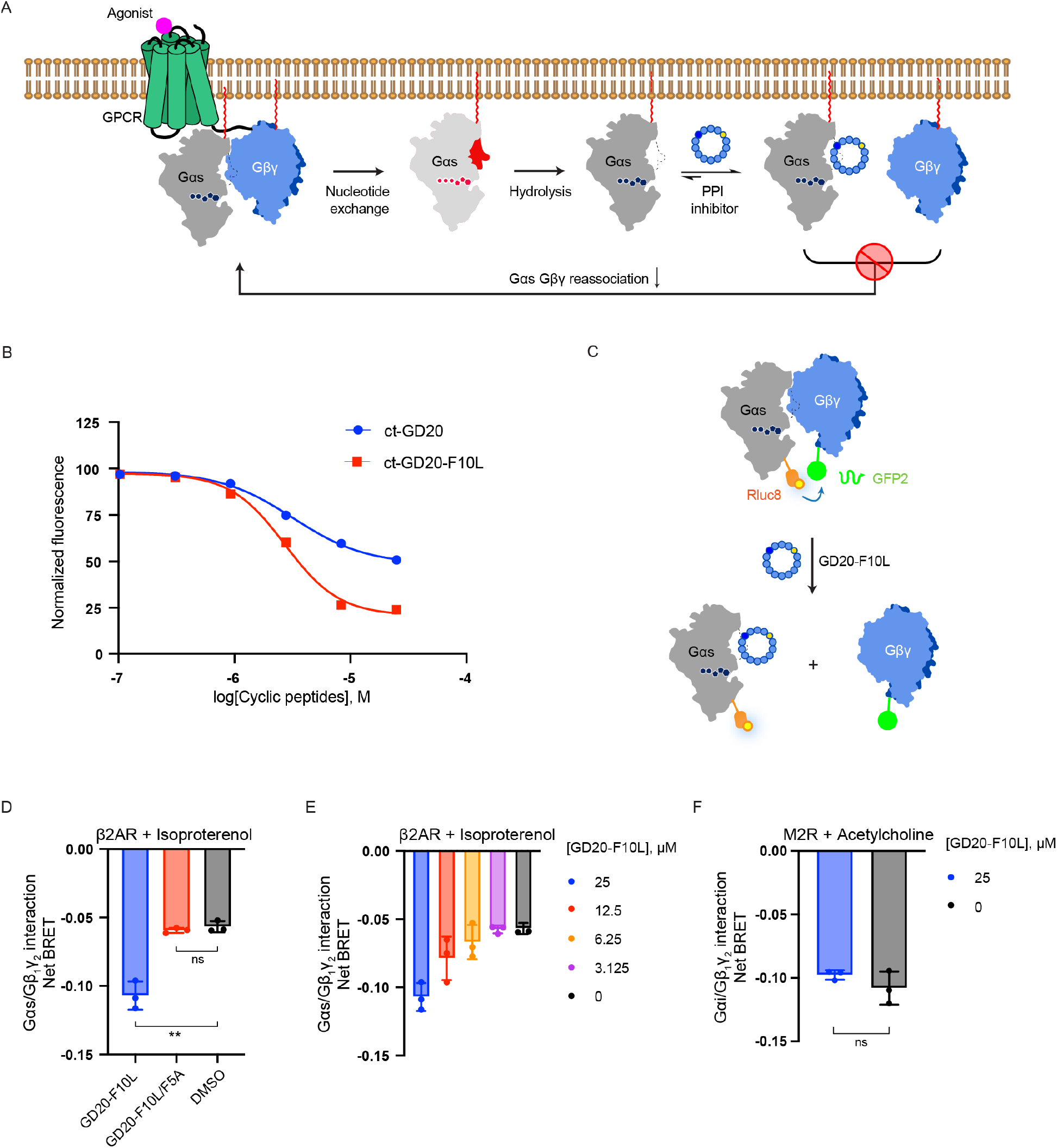
A cell permeable cyclic peptide GD20-F10L inhibits Gαs Gβγ reassociation in HEK293 cells. (A) Schematic representation of PPI inhibitors inhibiting Gαs Gβγ reassociation in HEK 239 cells. Gαs/Gβγ trimer is first dissociated by GPCR activation. PPI inhibitors captures the monomeric Gαs/GDP after Gαs/GTP hydrolysis, which prevents Gαs Gβγ reassociation. (B) CAPA cell permeability assay results for ct-GD20 (blue) and ct-GD20-F10L (red). Each point is the median ct-TAMRA fluorescence of 10,000 cells. The data were normalized using cells that were only treated with ct-TAMRA as 100% signal and cells that were not treated with any ct-compound as 0% signal. The data represent the mean ± SD of three independent replicates. (C) Schematic representation of GD20-F10L inhibiting the protein-protein interaction between GαsShort_Rluc and Gβ1/GFP2_γ2 in a BRET2 assay. (D) HEK293 cells transfected with β2AR, Gαs-RLuc8, Gβ1, and Gγ2-GFP2 were pretreated with 25 µM GD20-F10L, GD20-F10L/F5A or DMSO for 16hours. Gαs/Gβγ dissociation was measured by BRET2 signal reduction after 10 nM isoproterenol application. BRET2 signal was normalized to cells that were not treated with isoproterenol. The data represent the mean ± SD of three independent replicates. Two-tailed unpaired t-tests were performed and P < 0.05 was considered significant. *p < 0.05, **p < 0.005, ns > 0.05. (E) HEK293 cells transfected with β2AR, Gαs-RLuc8, Gβ1, and Gγ2-GFP2 were pretreated with various concentrations of GD20-F10L for 16hours. Gαs/Gβγ dissociation was measured by BRET2 signal reduction after 10 nM isoproterenol application. BRET2 signal was normalized to cells that were not treated with isoproterenol. The data represent the mean ± SD of three independent replicates. (F) HEK293 cells transfected with M2R, Gαi1-RLuc8, Gβ1, and Gγ2-GFP2 were pretreated with 25 µM GD20-F10L or DMSO for 16hours. Gαi1/Gβγ dissociation was measured by BRET2 signal reduction after 100 nM acetylcholine application. BRET2 signal was normalized to cells that were not treated with acetylcholine. The data represent the mean ± SD of three independent replicates. Two-tailed unpaired t-tests were performed and P < 0.05 was considered significant. *p < 0.05, **p < 0.005, ns > 0.05. See also Figure S6.

We first tested the cell permeability of GD20, as peptide-based chemical probes often suffer from poor cell permeability. Several G protein-specific linear peptides exhibit *in vitro* activities but have no reported cellular efficacy, likely due to their low cell permeability (Ja and Roberts, 2004; Johnston et al., 2005; Johnston et al., 2005; Johnston et al., 2006; Austin et al., 2008). In order to quantitatively evaluate the cell permeability of GD20, we used a recently developed HaloTag-based assay known as a chloroalkane penetration assay (CAPA) (Figure S6I) (Peraro et al., 2018). HeLa cells stably expressing HaloTag localized to the mitochondrial outer membrane were pulsed with chloroalkane-tagged molecules (ct-molecule), washed, chased with chloroalkane-tagged TAMRA fluorophore (ct-TAMRA), and finally analyzed by flow cytometry. A lower ct-TAMRA fluorescent signal indicates competition from a higher cytosolic concentration of ct-molecule. The carboxyl terminus of GD20 (G15) was conjugated with a chloroalkane tag to make ct-GD20 (Figure S6A). While ct-GD20 is cell permeable, a single substitution F10L significantly improved cell penetration (Figure 6B, see also Figure S6B and S6C). Cyclic peptide GD20-F10L retains a similar level of binding affinity for GDP-bound Gαs with a *K*_D_ value of 14.5 ± 0.4 nM (Figure S6E), and comparable biochemical activity and class-specificity against Gαs Gβγ PPI, with an IC_50_ of 14.0 ± 0.6 nM for Gαs and a 100-fold selectivity over Gαi (Figure S6F to S6H).

We next tested GD20-F10L in HEK 293 cells overexpressing both β2AR and Gαs/Gβγ. Rluc8 was inserted within a flexible loop region between the αB-αC helices of Gαs (Figure S6J) and GFP2 was inserted at the N-terminus of Gγ2 to capture Gαβγ heterotrimer interaction. A decrease in the bioluminescence resonance energy transfer (BRET2) signal between labeled G protein subunits can detect Gαβγ dissociation (Figure 6C) in live cells (Olsen et al., 2020). We examined Gαβγ trimer dissociation elicited by the GPCR agonist at various concentrations of cyclic peptides by monitoring the net BRET2 signal. In cells that were transiently transfected with a Gαs-coupled β2-adrenergic receptor (β2AR), GαsShort_Rluc8, Gβ1, and Gγ-GFP2, ISO application stimulated a basal net BRET response. Pretreatment with GD20-F10L induced a greater net BRET2 signal between Gαs and Gβγ. This effect was GD20-F10L dose dependent (Figure 6D and 6E). In comparison, the Gαs binding deficient mutant GD20-F10L/F5A failed to induce a larger BRET2 response (Figure 6D). To assess the specificity of GD20-F10L at the G protein level, we tested it against the Gαi/Gβγ interaction. HEK 293 cells transiently expressing Gαi-coupled muscarinic acetylcholine receptor M2 (M2R), Gαi1_Rluc8, Gβ1, and Gγ-GFP2 were challenged with M2R agonist, acetylcholine. Pretreatment with GD20-F10L did not induce a net BRET2 signal change between Gαi and Gβγ (Figure 6F). These data suggested that GD20-F10L can specifically capture the monomeric state of Gαs after G protein activation and interfere with Gαs/Gβγ reassociation

## DISCUSSION

GPCRs and G proteins comprise the largest family of signal transducing proteins in the human genome. Although approximately 35% of approved drugs target GPCRs, directly targeting the downstream integrator G proteins has the potential for broader efficacy via blocking converged pathways shared by multiple GPCRs (Bonacci et al., 2006; Gulati et al., 2018). However, there is a striking absence of drug-like chemical matter that specifically targets the Gα proteins in cells. Cyclic peptides bridge the chemical space between small molecules and biologics, and are therefore capable of recognizing shallow effector binding pockets at PPI interfaces while maintaining optimal pharmacological properties. This is demonstrated here by the development of Gαs selective cyclic peptide inhibitors GN13 and GD20, which specifically recognize the Gαs switch II/α3 pocket, the site where downstream effectors bind. Cyclization of the peptide sequence and introduction of a non-canonical amino acid (D-tyrosine) provide these Gαs inhibitors better cell permeability and chemical stability (Figure 6B, see also Figure S6D, Table S4 and S5), making them comparable to small molecule drugs. Moreover, in contrast with the complex cyclic peptide natural product YM-254890, our Gαs-binding cyclic peptides can be easily derivatized through side-chain substitutions. The high-resolution co-crystal structures that we obtained of Gαs with our cyclic peptides enable us to program the protein-inhibitor interaction for desired biological effects. This tunability is exemplified by two GD20 analogs, GD20-F10L and GD20-F5A, in which a single point substitution drastically changed the biochemical and pharmacological properties of a given cyclic peptide, providing opportunities for further optimization.

Both GN13 and GD20 bind at the switch II/α3 pocket in Gαs. This pocket is evolutionally conserved and is commonly used for effector binding, with subtle differences conferred by sequence variability between homologous Gα proteins and by binding of different nucleotides (Wall et al., 1995; Tesmer et al., 1997; Slep et al., 2001; Tesmer et al., 2005; Chen et al., 2005; Liu et al., 2019). Our extremely diverse chemical library along with both positive and negative selection enabled us to survey the sequence space of cyclic peptides and discover selective binders that capture specific, subtly different conformations of the switch II/α3 pocket. The resulting Gαs-cyclic peptide recognitions are highly class-specific and state-selective, allowing for a precise modulation of yin and yang of Gαs signaling.

Gαs is one of the most frequently mutated G proteins in human cancer. Hotspot mutations in Gαs (Q227) disrupt its GTPase activity, thereby locking Gαs in the GTP-bound active conformation (Zachary et al., 1990). We found that the cyclic peptide GN13 recognized this particular Gαs conformation and inhibited GTP- and GppNHp-bound Gαs Q227L mutant in the adenylyl cyclase activation assay (Figure S2G). To our knowledge, this is the first demonstration of the ligandability of oncogenic Gαs.

The inactive state inhibitors GD20 and GD20-F10L similarly provide lead molecules to probe GDP-bound Gαs and exemplify a new mode of pharmacological intervention in GPCR signaling. The cell-penetrating cyclic peptide GD20-F10L captures a flexible switch-II conformation that is only available when Gαs is in the GDP bound state. This molecular recognition could be useful for developing biosensors directly detecting Gαs/GDP in cells. For example, a fluorescently tagged GD20-F10L could potentially be used for tracking real-time translocation of endogenous Gαs following receptor activation and internalization, which bypasses the need of G protein overexpression or genetic modification (Maziarz et al., 2020; Olsen et al., 2020).

GD20-F10L also offers a new angle to study the role of Gβγ signaling during GPCR stimulation. Gαs-GD20-F10L interaction sterically occludes Gβγ binding to Gαs. After acute stimulation of a Gαs-coupled receptor (β2AR), GD20-F10L functions as a dual-effect G protein PPI inhibitor by sequestering monomeric Gαs and releasing Gβγ from the natural inhibition of Gαs/GDP. As a result, GD20-F10L co-treatment with the β2AR agonist, isoproterenol generates a higher Gβγ concentration, which is comparable to the Gβγ concentration following M2R (a Gαi-coupled receptor) activation (Figure 6D and 6E). It was hypothesized that the level of free Gβγ upon receptor activation confers receptor signaling specificity (Touhara and MacKinnon, 2018). Therefore, GD20-F10L could potentially provide a novel approach to elucidate or even rewire the downstream signaling of Gαs-coupled receptors via activating Gβγ dependent pathways. Last, rapid Gα Gβγ reassociation terminates canonical GPCR-dependent G protein signaling within seconds (Ghosh et al., 2017). However, the slow dissociating Gαs-GD20-F10L interaction (Figure S6E and Table S3) offers the opportunity to trap the inactive state Gαs for a longer time and consequently prolong one branch of GPCR signaling — the Gβγ heterodimer.

Our demonstration of the use of the RaPID cyclic peptide platform through both positive and negative selection steps provides proof of principle for a path to discovering other cell-permeable, class-specific and state-selective inhibitors of the remainder of the GTPase family. The state-selective Gαs inhibitors GN13 and GD20 also provide novel pharmacological strategies for understanding and modulating GPCR signaling.

### Limitations of the study

Although GN13 and GD20-F10L are strong binders to Gαs, with *K*_D_ values in the nanomolar range, their potencies are compromised in our cell experiments. This is likely due to the difficulty of competing tight protein-protein interactions in cell membranes and the relatively lower cell penetration of cyclic peptides. Optimizing cyclic peptides with non-canonical residues could potentially improve the potency and cell permeability of GN13 and GD20-F10L to overcome this limitation. Second, although both GN13 and GD20-F10L exhibit excellent G protein class-specificity for Gαs over Gαi, we have not performed binding experiments with other Gα protein families (e.g., Gq-family and G12/13-family). It will be of interest in the future to test the specificity of GN13 and GD20-F10L against purified Gq and G12/13 proteins. Last, we investigated GD20-F10L activity with two well-studied GPCRs, β2AR and M2R. GD20-F10L showed great efficacy for the Gαs-coupled receptor, β2AR, but it would be worthwhile to test more Gαs-coupled receptors with GD20-F10L to further explore the scope of its utility.

## ACKNOWLEDGMENTS

We would like to thank the staff at A.L.S. beamline 8.2.1; Drs. Xiaobo Wan and Xi Liu for assistance in X-ray data processing and structure refinement; ELIM Biopharmaceuticals for their help with cyclic peptide synthesis; Drs. Roderick Mackinnon, Koki Tohara, and Reid Olsen for their help with developing cell-based BRET assays. Dr. Roshanak Irannejad, Dr. Kaihua Zhang, Jack Stevenson, and Doug Wassarman for comments on the manuscript. This work was supported by the Howard Hughes Medical Institute (K.M.S.); NIH grants R1R01CA244550 (K.M.S.), DA010711, DA012864, and MH120212 (M.v.Z.); Japan Agency for Medical Research and Development (AMED), Platform Project for Supporting Drug Discovery and Life Science Research (Basis for Supporting Innovative Drug Discovery and Life Science Research) under JP19am0101090 and the Japan Society for the Promotion of Science (JSPS) Grant-in-Aid for Specially Promoted Research JP20H05618 (H.S.). S.A.D is a UCSF Discovery fellow and a UCSF Fletcher Jones Fellow.; Q.H. is a Damon Runyon Fellow supported by the Damon Runyon Cancer Research Foundation (DRG-[2229-15]).; Z.Z. is a Damon Runyon Fellow supported by the Damon Runyon Cancer Research Foundation (DRG-[2281-17]).

## AUTHOR CONTRIBUTIONS

K.M.S., and H.S. conceived the project; S.A.D., Q.H., R.G., K.M.S., and H.S. designed the experiments. R.G. and H. P. performed cyclic peptide selection using the RaPID system; S.A.D., R.G., H. P. and Z.Z. carried out the chemical synthesis of cyclic peptides; S.A.D., and Q.H. performed biochemical characterization of the cyclic peptides, including protein purification, the adenylyl cyclase activation assay, the FRET assay, the BLI assay; Q.H. crystallized the GN13/Gαs and GD20/Gαs complexes and determined the structures; S.A.D. and E.E.B performed the cell-based assays; S.A.D., and K.M.S wrote the manuscript with the contribution from other authors.

## REFERENCES

Adams, P.D., Afonine, P. v., Bunkóczi, G., Chen, V.B., Davis, I.W., Echols, N., Headd, J.J., Hung, L.W., Kapral, G.J., Grosse-Kunstleve, R.W., et al. (2010). PHENIX: A comprehensive Python-based system for macromolecular structure solution. Acta Crystallogr. D Biol. Crystallogr. 66, 213–221.

Alessi, D.R., and Sammler, E. (2018). LRRK2 kinase in Parkinson’s disease. Science 360, 36–37.

Austin, R.J., Ja, W.W., and Roberts, R.W. (2008). Evolution of Class-Specific Peptides Targeting a Hot Spot of the Gαs Subunit. J. Mol. Biol. 377, 1406–1418.

Bonacci, T.M., Mathews, J.L., Yuan, C., Lehmann, D.M., Malik, S., Wu, D., Font, J.L., Bidlack, J.M. and Smrcka, A.V., 2006. Differential targeting of Gßγ-subunit signaling with small molecules. Science 312, 443–446.

Canon, J., Rex, K., Saiki, A.Y., Mohr, C., Cooke, K., Bagal, D., Gaida, K., Holt, T., Knutson, C.G., Koppada, N., et al. (2019). The clinical KRAS(G12C) inhibitor AMG 510 drives anti-tumour immunity. Nature 575, 217–223.

Chen, Z., Singer, W.D., Sternweis, P.C., and Sprang, S.R. (2005). Structure of the p115RhoGEF rgRGS domain-Gα13/i1 chimera complex suggests convergent evolution of a GTPase activator. Nat. Struct. Mol. Biol. 12, 191–197.

Dougherty, P.G., Sahni, A., and Pei, D. (2019). Understanding Cell Penetration of Cyclic Peptides. Chem. Rev. 119, 10241–10287.

Dror, R.O., Mildorf, T.J., Hilger, D., Manglik, A., Borhani, D.W., Arlow, D.H., Philippsen, A., Villanueva, N., Yang, Z., Lerch, M.T., et al. (2015). Structural basis for nucleotide exchange in heterotrimeric G proteins. Science 348, 1361–1365.

Emsley, P., Lohkamp, B., Scott, W.G., and Cowtan, K. (2010). Features and development of Coot. Acta Crystallogr. D Biol. Crystallogr. 66, 486–501.

Evans, P. (2006). Scaling and assessment of data quality. In Acta Crystallogr. D Biol. Crystallogr., pp. 72–82.

Ghosh, P., Rangamani, P., and Kufareva, I. (2017). The GAPs, GEFs, GDI. and…now, GEMs: New kids on the heterotrimeric G protein signaling block. Cell Cycle 16, 607–612.

Gilman, A.G. (1987). G proteins: transducers of receptor-generated signals. Annu Rev Biochem. 56, 615–649.

Goto, Y., Katoh, T., and Suga, H. (2011). Flexizymes for genetic code reprogramming. Nat Protoc. 6, 779–790.

Gulati, S., Jin, H., Masuho, I., Orban, T., Cai, Y., Pardon, E., Martemyanov, K.A., Kiser, P.D., Stewart, P.L., Ford, C.P. and Steyaert, J., (2018). Targeting G protein-coupled receptor signaling at the G protein level with a selective nanobody inhibitor. Nat. Commun. 9, 1–15.

Hallin, J., Engstrom, L.D., Hargi, L., Calinisan, A., Aranda, R., Briere, D.M., Sudhakar, N., Bowcut, V., Baer, B.R., Ballard, J.A., et al. (2020). The KRASG12C inhibitor MRTX849 provides insight toward therapeutic susceptibility of KRAS-mutant cancers in mouse models and patients. Cancer Discov. 10, 54–71.

Hu, Q., and Shokat, K.M. (2018). Disease-Causing Mutations in the G Protein Gαs Subvert the Roles of GDP and GTP. Cell 173, 1254–1264.e11.

Ja, W.W., and Roberts, R.W. (2004). In vitro selection of state-specific peptide modulators of G protein signaling using mRNA display. Biochemistry 43, 9265–9275.

Ja, W.W., Wiser, O., Austin, R.J., Jan, L.Y., and Roberts, R.W. (2006). Turning G proteins on and off using peptide ligands. ACS Chem. Biol. 1, 570–574.

Johnston, C.A., Willard, F.S., Jezyk, M.R., Fredericks, Z., Bodor, E.T., Jones, M.B., Blaesius, R., Watts, V.J., Harden, T.K., Sondek, J., et al. (2005). Structure of Gαi1 bound to a GDP-selective peptide provides insight into guanine nucleotide exchange. Structure 13, 1069–1080.

Johnston, C.A., Ramer, J.K., Blaesius, R., Fredericks, Z., Watts, V.J., and Siderovski, D.P. (2005). A bifunctional Gαi /Gαs modulatory peptide that attenuates adenylyl cyclase activity. FEBS Letters. 579, 5746–5750

Johnston, C.A., Lobanova, E.S., Shavkunov, A.S., Low, J., Ramer, J.K., Blaesius, R., Fredericks, Z., Willard, F.S., Kuhlman, B., Arshavsky, V.Y. and Siderovski, D.P., (2006). Minimal Determinants for Binding Activated Gα from the Structure of a Gαi1− Peptide Dimer. Biochemistry 45, 11390–11400.

Kaur, H., Harris, P.W.R., Little, P.J., and Brimble, M.A. (2015). Total synthesis of the cyclic depsipeptide YM-280193, a platelet aggregation inhibitor. Org. Lett. 17, 492–495.

Lambright, D.G., Noel, J.P., Hammt Be, H.E., and Sigler, P.B. (1994). Structural determinants for activation of the alpha-subunit of a heterotrimeric G protein. Nature 369, 621–628.

Liu, X., Xu, X., Hilger, D., Aschauer, P., Tiemann, J.K., Du, Y., Liu, H., Hirata, K., Sun, X., Guixà-González, R. and Mathiesen, J.M., (2019). Structural insights into the process of GPCR-G protein complex formation. Cell 177, 1243–1251.

Maziarz, M., Park, J.C., Leyme, A., Marivin, A., Garcia-Lopez, A., Patel, P.P., and Garcia-Marcos, M. (2020). Revealing the Activity of Trimeric G-proteins in Live Cells with a Versatile Biosensor Design. Cell 182, 770–785.e16.

McCoy, A.J., Grosse-Kunstleve, R.W., Adams, P.D., Winn, M.D., Storoni, L.C., and Read, R.J. (2007). Phaser crystallographic software. J Appl Crystallogr. 40, 658–674.

Morimoto, J., Hayashi, Y., and Suga, H. (2012). Discovery of macrocyclic peptides armed with a mechanism-based warhead: Isoform-selective inhibition of human deacetylase SIRT2. Angew Chem Int Ed Engl. 51, 3423–3427.

Murakami, H., Saito, H., and Suga, H. (2003). A Versatile tRNA Aminoacylation Catalyst Based on RNA. Chem Biol. 10, 655–662.

Murakami, H., Ohta, A., Ashigai, H., and Suga, H. (2006). A highly flexible tRNA acylation method for non-natural polypeptide synthesis. Nat Methods. 3, 357–359.

Neklesa, T.K., Tae, H.S., Schneekloth, A.R., Stulberg, M.J., Corson, T.W., Sundberg, T.B., Raina, K., Holley, S.A., and Crews, C.M. (2011). Small-molecule hydrophobic tagging-induced degradation of HaloTag fusion proteins. Nat. Chem. Biol. 7, 538–543.

Nishimura, A., Kitano, K., Takasaki, J., Taniguchi, M., Mizuno, N., Tago, K., Hakoshima, T., Itoh, H., and Gilman, A.G. (2010). Structural basis for the specific inhibition of heterotrimeric Gq protein by a small molecule. Proc. Natl. Acad. Sci. U.S.A. 107, 13666–13671.

O’Hayre, M., Vázquez-Prado, J., Kufareva, I., Stawiski, E.W., Handel, T.M., Seshagiri, S., and Gutkind, J.S. (2013). The emerging mutational landscape of G proteins and G-protein-coupled receptors in cancer. Nat. Rev. Cancer. 13, 412–424.

Olsen, R.H.J., DiBerto, J.F., English, J.G., Glaudin, A.M., Krumm, B.E., Slocum, S.T., Che, T., Gavin, A.C., McCorvy, J.D., Roth, B.L., et al. (2020). TRUPATH, an open-source biosensor platform for interrogating the GPCR transducerome. Nat. Chem. Biol. 16, 841–849.

Otwinowski, Z., and Minor, W. (1997). [20] Processing of X-ray diffraction data collected in oscillation mode. In Methods in enzymology (Vol. 276, pp. 307–326). Academic press.

Passioura, T., and Suga, H. (2017). A RaPID way to discover nonstandard macrocyclic peptide modulators of drug targets. Chem. Commun. (Camb.). 53, 1931–1940.

Peraro, L., Deprey, K.L., Moser, M.K., Zou, Z., Ball, H.L., Levine, B., and Kritzer, J.A. (2018). Cell Penetration Profiling Using the Chloroalkane Penetration Assay. J. Am. Chem. Soc. 140, 11360–11369.

Prior, I.A., Lewis, P.D., and Mattos, C. (2012). A comprehensive survey of ras mutations in cancer. Cancer Res. 72, 2457–2467.

Ramaswamy, K., Saito, H., Murakami, H., Shiba, K., and Suga, H. (2004). Designer ribozymes: Programming the tRNA specificity into flexizyme. J Am Chem Soc. 126, 11454–11455.

Slep, K.C., Kercher, M.A., Hek, W., Cowan, C.W., Wenselk, T.G., and Sigler, P.B. (2001). Structural determinants for regulation of phosphodiesterase by a G protein at 2.0 Å. Nature 409, 1071–1077.

Sohrabi, C., Foster, A., and Tavassoli, A. (2020). Methods for generating and screening libraries of genetically encoded cyclic peptides in drug discovery. Nat. Rev. Chem. 4, 90–101.

Stallaert, W., van der Westhuizen, E.T., Schönegge, A.M., Plouffe, B., Hogue, M., Lukashova, V., Inoue, A., Ishida, S., Aoki, J., le Gouill, C., et al. (2017). Purinergic receptor transactivation by the β2-adrenergic receptor increases intracellular Ca2+ in nonexcitable cells. Mol. Pharmacol. 91, 533–544.

Sunahara, R.K., Tesmer, J.J., Gilman, A.G. and Sprang, S.R., (1997). Crystal structure of the adenylyl cyclase activator Gsα. Science 278, 1943–1947.

Takasaki, J., Saito, T., Taniguchi, M., Kawasaki, T., Moritani, Y., Hayashi, K., and Kobori, M. (2004). A novel Gαq/11-selective inhibitor. J. Biol. Chem. 279, 47438–47445.

Tesmer, J.J.G., Sunahara, R.K., Gilman, A.G., and Sprang, S.R. (1997). Crystal Structure of the Catalytic Domains of Adenylyl Cyclase in a Complex with Gsα GTPγS. Science 278, 1907–1916

Tesmer, V.M., Kawano, T., Shankaranarayanan, A., Kozasa, T. and Tesmer, J.J., (2005). Snapshot of activated G proteins at the membrane: the Gαq-GRK2-Gßγ complex. Science 310, 1686–1690.

Touhara, K.K. and MacKinnon, R., (2018). Molecular basis of signaling specificity between GIRK channels and GPCRs. Elife 7, e42908.

Wall, M.A., Coleman, D.E., Lee, E., IAiguez-Lluhi, J.A., Posner, B.A., Gilman, A.G., and Ft Sprang, S. (1995). The Structure of the G Protein Heterotrimer Giα1β1γ2. Cell 83, 1047–1058

Weis, W.I., and Kobilka, B.K. (2018). The Molecular Basis of G Protein-Coupled Receptor Activation. Annu. Rev. Biochem. 87, 897–919.

Wilson, G.R., Sim, J.C.H., McLean, C., Giannandrea, M., Galea, C.A., Riseley, J.R., Stephenson, S.E.M., Fitzpatrick, E., Haas, S.A., Pope, K., et al. (2014). Mutations in RAB39B cause X-linked intellectual disability and early-onset parkinson disease with α-synuclein pathology. Am. J. Hum. Genet. 95, 729–735.

Winn, M.D., Ballard, C.C., Cowtan, K.D., Dodson, E.J., Emsley, P., Evans, P.R., Keegan, R.M., Krissinel, E.B., Leslie, A.G.W., McCoy, A., et al. (2011). Overview of the CCP4 suite and current developments. Acta Crystallogr. D Biol. Crystallogr. 67, 235–242.

Xiao, H., Murakami, H., Suga, H., and Ferré-D’Amaré, A.R. (2008). Structural basis of specific tRNA aminoacylation by a small in vitro selected ribozyme. Nature 454, 358–361.

Xiong, X.F., Zhang, H., Underwood, C.R., Harpsøe, K., Gardella, T.J., Wöldike, M.F., Mannstadt, M., Gloriam, D.E., Bräuner-Osborne, H., and Strømgaard, K. (2016). Total synthesis and structure-activity relationship studies of a series of selective G protein inhibitors. Nat Chem. 8, 1035–1041.

Yamagishi, Y., Shoji, I., Miyagawa, S., Kawakami, T., Katoh, T., Goto, Y., and Suga, H. (2011). Natural product-like macrocyclic N-methyl-peptide inhibitors against a ubiquitin ligase uncovered from a ribosome-expressed de novo library. Chem Biol. 18, 1562–1570.

Zachary, I., Masters, S.B. and Bourne, H.R., (1990). Increased mitogenic responsiveness of Swiss 3T3 cells expressing constitutively active Gsα. Biochem. Biophys. Res. Commun. 168, 1184–1193.

Zhang, H., Xiong, X.F., Boesgaard, M.W., Underwood, C.R., Bräuner-Osborne, H., and Strømgaard, K. (2017). Structure–Activity Relationship Studies of the Cyclic Depsipeptide Natural Product YM-254890, Targeting the Gq Protein. ChemMedChem 12, 830–834.

