## Supplementary Materials for "State-selective Modulation of Heterotrimeric Gαs Signaling with Macrocyclic Peptides"

This PDF file includes:

Materials and Methods
Figs. S1 to S6
Tables S1 to S5

### RESOURCE AVAILABILITY

#### *Lead Contact*

#### *Materials Availability*

All unique reagents generated in this study are available from the lead contact but we may require a completed Materials Transfer Agreement if there is potential for commercial application.

#### *Data and code availability*

Data Resources: X-ray Crystallography data have been deposited at Protein Data Bank (PDB) and are publicly available as of the date of publication. The accession number for the crystal structure of GppNHp-bound Gαs in complex with the cyclic peptide inhibitor GN13 reported in this paper is PDB: 7BPH. The accession number for the crystal structure of GDP-bound Gαs in complex with the cyclic peptide inhibitor GD20 reported in this paper is PDB: 7E5E.

### EXPERIMENTAL MODEL AND SUBJECT DETAILS

#### *Cell lines*

HeLa cells stably expressing the Halo-Tag-GFP-Mito construct were provided by the Kritzer lab (Peraro et al., 2018). Wild-type HEK293, GNAS KO HEK293 were provided by the Inoue lab. These cells are female in origin. Wild-type HEK293, GNAS KO HEK293 and HeLa cells were cultured at 37 °C, 5% $CO_2$ in DMEM (Thermo Fisher Scientific, Cat# 11995073) supplemented with 10% heat-inactivated FBS (AxeniaBiologix).

WT Gαs, all the mutants of Gαs, the C1 domain (residues 442-658, VC1) of human ADCY5 (adenylyl cyclase V) and the C2 domain (residues 871-1082, IIC2) of human ADCY2 (adenylyl cyclase II) were overexpressed in *Escherichia coli* BL21(DE3) cultured in Terrific Broth (TB) Medium. Human GNB1 (Gβ1) and GNG2 (Gγ2) were co-expressed in Sf9 insect cells cultured in Sf-900 III SFM medium at 28 °C.

### METHOD DETAILS

#### *Protein expression and purification*

##### *Proteins used in the adenylyl cyclase assay, the radioactivity assay, and the steady state GTPase asssy*

The wild-type and S275L mutant of Gαs, C2 domain of human ADCY2, C1 domain of human ADCY5, and human Gβ1/Gγ2(C68S) complex used in the adenylyl cyclase activity assay were cloned, expressed and purified as described (Hu and Shokat, 2018).

*Gαs used in the RaPID selection*

The gene encoding residues 7-380 of the short isoform of human Gαs (*GNAS*, accession number in PubMed: NP_536351) with an Avi tag and a TEV cleavage site at its N-terminus was cloned into the multiple cloning site 1 of the pETDuet vector. The resulting protein sequence is as follows:

MGSSHHHHHHSGMSGLNDIFEAQKIEWHESSGENLYFQGMSKTEDQRNEEKAQREA
NKKIEKQLQKDKQVYRATHRLLLLGAGESGKSTIVKQMRILHVNGFNGDSEKATKVQDI
KNNLKEAIETIVAAMSNLVPPVELANPENQFRVDYILSVMNVPDFDFPPEFYEHAKALW
EDEGVRACYERSNEYQLIDCAQYFLDKIDVIKQADYVPSDQDLLRCRVLTSGIFETKFQ
VDKVNFHMFDVGGQRDERRKWIQCFNDVTAIIFVVASSSYNMVIREDNQTNRLQEALN
LFKSIWNNRWLRTISVILFLNKQDLLAEKVLAGKSKIEDYFPEFARYTTPEDATPEPGED
PRVTRAKYFIRDEFLRISTASGDGRHYCYPHFTCAVDTENIRRVFNDCRDIIQRMHLRQ
YELL

In the same pETDuet plasmid, the gene encoding BirA (accession number in PubMed: NP_418404.1) was inserted between NdeI and XhoI sites of the multiple cloning site 2. This plasmid was transformed into *Escherichia coli* BL21(DE3). The transformed cells were grown in TB medium supplemented with 50 μg/mL carbenicillin at 37 °C until OD600 reached 0.5, and then cooled to 22 °C followed by addition of 40 μM β-D-thiogalactopyranoside. After overnight incubation, 50 μM biotin was added into the culture for 2 hours. The cells were harvested by centrifugation, resuspended in lysis buffer (150 mM NaCl, 25 mM Tris 8.0, 1 mM $MgCl_2$, 250 μM biotin, protease inhibitor, and then lysed by a microfluidizer. The cell lysate was centrifuged at 14000 g for 1 hour at 4 °C. The supernatant was incubated with TALON Resin at 4 °C for 2 hours, then the resin was washed by 500 mM NaCl, 25 mM Tris 8.0, 1 mM $MgCl_2$ and 5 mM imidazole 8.0. Gαs was eluted by 25 mM Tris 8.0, 1 mM $MgCl_2$, 250 mM imidazole 8.0, 10% glycerol and 0.1 mM GDP. After adding 5 mM Dithiothreitol (DTT), the eluate was loaded onto a Source-15Q column. Gαs was eluted by a linear gradient from 100% IEC buffer A (25 mM Tris 8.0, 1 mM $MgCl_2$) to 40% IEC Buffer B (25 mM Tris 8.0, 1 M NaCl, 1 mM $MgCl_2$). The peak fractions were pooled and supplemented with 5 mM DTT. One half of peak fractions was mixed with equal volume of GppNHp exchange buffer (150 mM NaCl, 25 mM HEPES 8.0, 2 mM EDTA, 2 mM GppNHp, 5 mM DTT) for 2 hours, followed by addition of 5 mM $MgCl_2$. GppNHp-bound Gαs and GDP-bound Gαs were concentrated and purified by gel filtration (Superdex 200 increase, 10/30) with SEC buffer (150 mM NaCl, 20 mM HEPES 8.0, 5 mM $MgCl_2$ and 1 mM EDTA-Na 8.0). The peak fractions were pooled and concentrated for biochemical assay.

*WT Gαs, Gαs S275L mutant and full-length Gαi used in the TR-FRET assay and the bio-layer interferometry assay*

The gene of residues 7-380 of the short isoform of human Gαs (GNAS, accession number in PubMed: NP_536351) with a stop codon at its end was cloned into the NdeI/XhoI site of a modified pET15b vector, in which a Drice cleavage site (AspGluValAsp↓Ala) and an Avi tag were inserted at the N-terminus. The resulting protein sequence after Drice protease cleavage is as follows:

AHMGLNDIFEAQKIEWHESKTEDQRNEEKAQREANKKIEKQLQKDKQVYRATHRLLLL
GAGESGKSTIVKQMRILHVNGFNGDSEKATKVQDIKNNLKEAIETIVAAMSNLVPPVEL
ANPENQFRVDYILSVMNVPDFDFPPEFYEHAKALWEDEGVRACYERSNEYQLIDCAQ
YFLDKIDVIKQADYVPSDQDLLRCRVLTSGIFETKFQVDKVNFHMFDVGGQRDERRKW
IQCFNDVTAIIFVVASSSYNMVIREDNQTNRLQEALNLFKSIWNNRWLRTISVILFLNKQD
LLAEKVLAGKSKIEDYFPEFARYTTPEDATPEPGEDPRVTRAKYFIRDEFLRISTASGDG
RHYCYPHFTCAVDTENIRRVFNDCRDIIQRMHLRQYELL

The AviTagged Gαs S275L mutant plasmid was constructed using quick-change mutagenesis from the AviTagged WT Gαs plasmid. The resulting protein sequence after Drice protease cleavage is as follows:
AHMGLNDIFEAQKIEWHESKTEDQRNEEKAQREANKKIEKQLQKDKQVYRATHRLLLL
GAGESGKSTIVKQMRILHVNGFNGDSEKATKVQDIKNNLKEAIETIVAAMSNLVPPVEL
ANPENQFRVDYILSVMNVPDFDFPPEFYEHAKALWEDEGVRACYERSNEYQLIDCAQ
YFLDKIDVIKQADYVPSDQDLLRCRVLTSGIFETKFQVDKVNFHMFDVGGQRDERRKW
IQCFNDVTAIIFVVASSSYNMVIREDNQTNRLQEALNLFKLIWNNRWLRTISVILFLNKQD
LLAEKVLAGKSKIEDYFPEFARYTTPEDATPEPGEDPRVTRAKYFIRDEFLRISTASGDG
RHYCYPHFTCAVDTENIRRVFNDCRDIIQRMHLRQYELL

The gene of residues 2-354 of the short isoform of human Gαi1 (GNAI1, accession number in PubMed: NP_002060.4) with a stop codon at its end was cloned into the NdeI/XhoI site of a modified pET15b vector, in which a Drice cleavage site (AspGluValAsp↓Ala) and an Avi tag were inserted at the N-terminus. The resulting protein sequence after Drice protease cleavage is as follows:
AHMGLNDIFEAQKIEWHEGCTLSAEDKAAVERSKMIDRNLREDGEKAAREVKLLLLGA
GESGKSTIVKQMKIIHEAGYSEEECKQYKAVVYSNTIQSIIAIIRAMGRLKIDFGDSARAD
DARQLFVLAGAAEEGFMTAELAGVIKRLWKDSGVQACFNRSREYQLNDSAAYYLNDL
DRIAQPNYIPTQQDVLRTRVKTTGIVETHFTFKDLHFKMFDVGGQRSERKKWIHCFEG
VTAIIFCVALSDYDLVLAEDEEMNRMHESMKLFDSICNNKWFTDTSIILFLNKKDLFEEKI
KKSPLTICYPEYAGSNTYEEAAAYIQCQFEDLNKRKDTKEIYTHFTCATDTKNVQFVFD
AVTDVIIKNNLKDCGLF

The above-mentioned plasmids were transformed into *Escherichia coli* BL21(DE3), respectively. The transformed cells were grown in TB medium supplemented with 50 μg/mL carbenicillin at 37 °C until OD600 reached 0.4, and then cooled to 22 °C followed by addition of 100 μM IPTG. After overnight incubation, the cells were harvested by centrifugation, resuspended in lysis buffer (150 mM NaCl, 25 mM Tris 8.0, 1 mM $MgCl_2$, protease inhibitor cocktail), and then lysed by a microfluidizer. The cell lysate was centrifuged at 14000 g for 1 hour at 4 °C. The supernatant was incubated with TALON resin at 4 °C for 1 hour, then the resin was washed by 500 mM NaCl, 25 mM Tris 8.0, 1 mM $MgCl_2$ and 5 mM imidazole 8.0. G protein was eluted by 25 mM Tris 8.0, 1 mM $MgCl_2$, 250 mM imidazole 8.0, 10% glycerol and 0.1 mM GDP. After adding 5 mM Dithiothreitol (DTT), the eluate was incubated with Drice protease at 4 °C overnight to remove the hexahistidine tag. Purified BirA (A gift from the Wells lab) and biotin were added at 4 °C until LC-MS showed complete biotinylation. G protein was loaded onto a Source-15Q column and eluted by a linear gradient from 100% IEC buffer A (25 mM Tris 8.0, 1 mM

$MgCl_2$) to 40% IEC Buffer B (25 mM Tris 8.0, 1 M NaCl, 1 mM $MgCl_2$). The peak fractions were pooled, nucleotide exchanged, and supplemented with 5 mM DTT and 0.1 mM nucleotide, and then concentrated and purified by gel filtration (Superdex 200 increase, 10/30) with SEC buffer (150 mM NaCl, 20 mM HEPES 8.0, 5 mM $MgCl_2$ and 1 mM EDTA-Na 8.0). The peak fractions were pooled and concentrated for biochemical assay.

*RaPID Selection*

Selections were performed with thioether-macrocyclic peptide library against biotinylated Gαs. Thioether-macrocyclic peptide libraries were constructed with N-chloroacetyl-D-tyrosine (ClAc$^{D}$Tyr) as an initiator by using the flexible in vitro translation (FIT) system (Goto et al., 2011). The mRNA libraries, ClAc$^{D}$Tyr-tRNA$^{fMet}_{CAU}$ were prepared as reported (Yamagishi et al., 2011). The mRNA library corresponding for the thioether-macrocyclic peptide library was designed to have an AUG initiator codon to incorporate N-chloroacetyl-D-tyrosine (ClAc$^{D}$Tyr), followed by 8–12 NNK random codons (N = G, C, A or U; K = G or U) to code random proteinogenic amino acids, and then a fixed downstream UGC codon to assign Cys. After *in vitro* translation, a thioether bond formed spontaneously between the N-terminal ClAc group of the initiator $^{D}$Tyr residue and the sulfhydryl group of a downstream Cys residue.

In the first round of selection, the initial cyclic peptide library was formed by adding puromycin ligated mRNA library (225 pmol) to a 150 µL scale flexible *in vitro* translation system, in the presence of 30 µM of ClAc$^{D}$Tyr-tRNA$^{fMet}_{CAU}$. The translation was performed 37 °C for 30 min, followed by an extra incubation at 25 °C for 12 min. After an addition of 15 µL of 200 mM EDTA (pH 8.0) solution, the reaction solution was incubated at 37 °C for 30 min to facilitate cyclization. Then the library was reversed transcribed by M-MLV reverse transcriptase at 42 °C for 1 hour and subject to pre-washed Sephadex G-25 columns to remove salts. The desalted solution of peptide–mRNA/cDNA was applied to Gαs (positive selection state)-immobilized Dynabeads M280 streptavidin magnetic beads and rotated at 4 °C for 1 hour in selection buffer (25 mM HEPES pH 7.5, 150 mM NaCl, 1 mM $MgCl_2$ and 0.05% Tween 20) containing 0.5 mM corresponding nucleotide and 0.1% acetylated BSA. Bead amounts were chosen that the final concentration of Gαs protein was 200 nM. This process is referred to as positive selection. The selected peptide–mRNA/cDNAs were isolated from the beads by incubating in 1xPCR reaction buffer heated at 95 °C for 5 min. The amount of eluted cDNAs was measured by quantitative PCR. The remaining cDNAs were amplified by PCR, purified and transcribed into mRNAs as a library for the next round of selection.

In the subsequent rounds of selection, ligated mRNA from previous round (7.5 pmol) was added to a 5 µL scale reprogrammed *in vitro* translation system. This was incubated at 37 °C for 30 min and at 25 °C for 12 min. Then 1 µL of 100 mM EDTA (pH 8.0) was added and incubated at 37 °C for 30 min. After reverse transcription and subject to pre-washed Sephadex G-25 columns to remove salts, negative selection was performed by adding the desalted solution of peptide–mRNA/cDNA to Gαs (negative selection state)-immobilized Dynabeads M280 streptavidin magnetic beads and rotated at 4 °C for 30 min in selection buffer containing 0.1% acetylated BSA. This process was repeated several times by removing the supernatant to fresh beads immobilized with Gαs (negative selection state). The supernatant from the last negative selection was then added to beads immobilized with the positive selection state of Gαs (final conc. 200nM) and rotated at 4 °C for 30 min in selection buffer containing 0.5mM corresponding nucleotide and 0.1% acetylated BSA. As described in the first round of selection, the cDNA was quantified with qPCR, amplified with PCR, transcribed and ligated to puromycin. The subsequent selection was repeated for several rounds until a significant enrichment of cDNA was observed for positive selection state. The recovered cDNA was then identified by next generation sequencing (Miseq, Illumina).

#### *Comparison selection*

In comparison selection, ligated mRNA (7.5 pmol) from last round selection was added to a 5 μL scale reprogrammed in *vitro* translation system. After translation, cyclization, reverse transcription and prewashed with Sephadex G-25 columns, the desalted solution of peptide–mRNA/cDNA library was split equally into three fractions, and perform three paralleled selections with the same amount of blank, GDP-bound Gαs-immobilized or GppNHp-bound Gαs-immobilized Dynabeads M280 streptavidin magnetic beads, individually. For each of the paralleled selections, the beads were rotate at 4 °C for 30 min, washed three times with selection buffer. The remaining cDNAs were then eluted from the beads, quantified by qPCR, followed by Miseq sequencing. Finally, identified sequences from each paralleled selection were compared by normalization of Miseq abundance of the sequence with the qPCR reads of the paralleled selection.

#### *Bio-layer interferometry (BLI)*

BLI experiments were performed using an OctetRED384 instrument from ForteBio. All experiments were performed at 25 °C using BLI buffer (10 mM HEPES pH 7.4, 150 mM NaCl, 1mM $MgCl_2$, 0.05% Tween-20, 0.1% DMSO, 0.2mM GppNHp or GDP). Cyclic peptides were diluted to a series of concentrations in BLI buffer plus 10 μM Biotin. Assays were conducted in Greiner 384well, black, flat bottom polypropylene plates containing the protein solutions, BLI buffer plus 10 μM Biotin for dissociation, and serial dilutions of cyclic peptides to be tested.

Biotinylated proteins were immobilized on Streptavidin biosensors by dipping sensors into plate wells containing protein solutions at a concentration of 100 - 150 nM. Protein loading is around 2-3 nm. Sensors loaded with proteins were moved and dipped into wells with BLI buffer plus 10 μM Biotin to block unlabeled Streptavidin. Association–dissociation cycles of compounds were started by moving and dipping sensors to cyclic peptides dilutions and BLI buffer plus 10 μM Biotin wells alternatively. Association and dissociation times were carefully determined to ensure full association and dissociation.

Raw kinetic data collected were processed with the Data Analysis software provided by the manufacturer using single reference subtraction in which buffer-only reference was subtracted (For GN13 analysis). Because GD20 analogs have a low level of background binding, we used a double reference subtraction (buffer-only reference and non-protein-loading reference) method to calculate their kinetics values. The resulting data were analyzed based on a 1:1 binding model from which $k_{on}$ and $k_{off}$ values were obtained and then $K_d$ values were calculated.

#### *Adenylyl cyclase activity assay*

*Cyclic peptides dose dependent inhibition (Figure 2D).*
Cyclic peptides GN1, GN3, GN6, GN7, GN8, GN10, GN11, GN13, GN15 (4 mM stock in DMSO) were diluted to 4X stocks with a series of concentrations (0, 1.56, 3.12, 6.25, 12.5, 25, 50, 100 µM) in reaction buffer (1x PBS 7.4, 0.1% BSA). WT Gαs at a concentration of 8.5 mg/mL (about 190 µM) in 20 mM HEPES 8.0, 150 mM NaCl, 5 mM $MgCl_2$, 1 mM EDTA-Na 8.0 was diluted to 0.5 µM in dilution buffer (1x PBS 7.4, 0.1% BSA, 1 mM EDTA-Na 8.0, 2 mM DTT, 0.1mM $MgCl_2$) plus 1mM GppNHp. After incubation at room temperature for 1 hour to allow nucleotide exchange, 2.5 µL of Gαs dilution was mixed with 1 µL $MgCl_2$ stock (20 mM $MgCl_2$, 1x PBS 7.4, 0.1% BSA) in an OptiPlate-384, White Opaque 384-well Microplate to lock Gαs in GppNHp-bound state. 2 µL of adenylyl cyclase stock (2 µM VC1, 2 nM IIC2, 150 µM FSK, 1x PBS 7.4, 0.1% BSA) was added, followed by addition of 2.5 µL 4X cyclic peptides stock. Reaction mixture was further incubated at room temperature for 2 hours and placed on ice for 5 minutes. cAMP production was initiated by addition of 2 µL of ATP stock (1 mM ATP, 1x PBS 7.4, 0.1% BSA). The reaction was carried out at 30 °C for 10 minutes in a PCR machine and stopped by heating at 95 °C for 3 minutes. The cAMP concentrations were measured by the LANCE Ultra cAMP kit.

*GN13 inhibition of Gαs proteins at various concentrations (Figure 3H).*
WT Gαs and S275L mutant at a concentration of 8.5 mg/mL (about 190 µM) in 20 mM HEPES 8.0, 150 mM NaCl, 5 mM $MgCl_2$, 1 mM EDTA-Na 8.0 were diluted to a series of concentrations (4 µM, 1.33 µM, 0.44 µM, 0.15 µM, 49.4 nM, 16.5 nM, 5.5 nM, 0 nM) in dilution buffer (1x PBS 7.4, 0.1% BSA, 1 mM EDTA-Na 8.0, 2 mM DTT, 0.1mM $MgCl_2$) plus 1mM GppNHp. After incubation at room temperature for 1 hour to allow nucleotide exchange, 2.5 µL of each sample was then mixed with 1µL of $MgCl_2$ stock (20 mM $MgCl_2$, 1x PBS 7.4, 0.1% BSA) in an OptiPlate-384, White Opaque 384-well Microplate. 2 µL of adenylyl cyclase/Gβγ stock (2 µM VC1, 2 nM IIC2, 150 µM FSK, 1x PBS 7.4, 0.1% BSA, 10 µM Gβ1/γ2(C68S)) was added, followed by addition of 2.5 µL 25 µM GN13 stock in 1x PBS 7.4, 0.1% BSA. Reaction mixture was further incubated at room temperature for 2 hours and placed on ice for 5 minutes. cAMP production was initiated by addition of 2 µL of ATP stock (1 mM ATP, 1x PBS 7.4, 0.1% BSA). The reaction was carried out at 30 °C for 10 minutes in a PCR machine and stopped by heating at 95 °C for 3 minutes. The cAMP concentrations were measured by the LANCE Ultra cAMP kit.

*GN13 inhibition of Gαs proteins in HEK293 cell membranes (Figure 2F and 3I).*
a. Cell membrane preparation: HEK293cells, GNAS KO HEK293 cells were plated two day before transfection at a density of 1M cells per 10cm plate. One plate of GNAS KO HEK293 cells was transfected with 4 µg of GNAS WT or GNAS S275L plasmids. After overnight transfection, cells were lifted with TypLE, washed, resuspended in stimulation buffer (1X PBS, protease inhibitor cocktail, 5 mM $MgCl_2$). Cell membranes were disrupted

by using the Dounce homogenizer for 25 strokes. Nuclei and unbroken cells were removed by centrifugation for 5 min at 500 g. The supernatant suspension was carefully removed and centrifuged for 30 min at 45K g. Cell membranes were suspended in stimulation buffer. The protein concentrations were measured using BCA, and were normalized to 750 µg/mL. A final concentration of 0.1% BSA was added into the cell membrane suspension. b: Adenylyl cyclase activity assay in cell membranes: 600 µL of cell membrane suspension was mixed with 60 µL of GTP/GDP stock (stock concentration: 10 mM/1 mM). 5.5 µL of the mixture from last step was mixed with 5.5 µL of GN13 and incubated at room temperature. After 2 hours, membrane/cyclic peptide mixture was transferred on ice for 5 minutes, followed by the addition of 2 µL of IBMX/ISO/ATP or IBMX/DMSO/ATP stock (5 mM IBMX, 0.2 mM ISO or DMSO, 2.5 mM ATP in stimulation buffer with 0.1% BSA). The reaction was carried out at 30 °C for 30 minutes in a PCR machine and stopped by heating at 95 °C for 3 minutes. The cAMP concentrations were measured by the LANCE Ultra cAMP kit.

*cAMP concentrations measurement by the LANCE Ultra cAMP kit.*
A cAMP standard curve was generated in the same plate using the 50 µM cAMP standard in the kit. Before the measurement, the samples were diluted by stimulation buffer (1x PBS 7.4, 0.1% BSA) to 1/60, 1/120, 1/240 or 1/480 to make sure the cAMP concentrations were in the dynamic range of the cAMP standard curve. 10 µL of each diluted sample was mixed with 5 µL of 4X Eu-cAMP tracer and 5 µL of 4X ULight-anti-cAMP in a white, opaque Optiplate-384 microplate, incubated for 1 hour at room temperature, and the time-resolved fluorescence resonance energy transfer (TR-FRET) signals were read on a Spark 20M plate reader. The cAMP standard curve was fitted by the software GraphPad Prism using the following equation in which ‘‘Y’’ is the TR-FRET signal and ‘‘X’’ is the log of cAMP standard concentration (M):

Y = Bottom + (Top-Bottom)/(1 + 10^ ((LogIC50-X)* HillSlope))

After obtained the values of the four parameters ‘‘Bottom’’, ‘‘Top’’, ‘‘LogIC50’’ and ‘‘HillSlope’’, we used this equation to convert the TR-FRET signals of the samples into cAMP production values. The cyclic peptides dose dependent inhibition curves were fitted by the following equation to calculate the IC50 of each cyclic peptide:

Y = Bottom + (Top-Bottom)/(1 + 10^ ((LogIC50-X)* HillSlope)),

in which ‘‘Y’’ is the cAMP production value, ‘‘X’’ is the log of cyclic peptide concentration (M).

*FRET based Gαs/adenylyl cyclase interaction assay*

Cyclic peptides GN13 (4 mM stock in DMSO) were diluted to 5X stocks with a series of concentrations (0, 0.0034, 0.0102, 0.0305, 0.0914, 0.274, 0.823, 2.47, 7.41, 22.2, 66.7, 200 µM) in 1X PBS 7.4, 0.1% BSA, 2 mM DTT, 2 mM $MgCl_2$. WT Gαs and Gαs S275L mutant at a concentration of 4.6 mg/mL (about 100 µM) in 20 mM HEPES 8.0, 150 mM NaCl, 5 mM $MgCl_2$ were diluted to 4 µM in EDTA GppNHp buffer (1x PBS 7.4, 0.1% BSA, 2 mM EDTA-Na 8.0, 2 mM DTT, 0.1mM $MgCl_2$, 1mM GppNHp). After incubation at room temperature for 1 hour to allow nucleotide exchange, Gαs dilutions were mixed with equal volume of $MgCl_2$ stock (3.8 mM $MgCl_2$, 1x PBS 7.4, 0.1% BSA, 2mM DTT) to lock Gαs in GppNHp-bound state. GppNHp-bound Gαs proteins were then diluted to 500 nM (5X stocks) in 1X PBS 7.4, 0.1% BSA, 2 mM DTT, 2 mM $MgCl_2$ plus 0.5 mM GppNHp. In an OptiPlate-384 White Opaque 384-well Microplate, 5X Gαs proteins were mixed with 5X GN13 serial dilution stocks, 5X streptavidin XL665 stock (125 nM), 5X adenylyl cyclase stock (VC1: 100 nM, IIC2: 200 nM, FSK 0.5mM) and 5X anti-6His-Tb cryptate stock (0.26 μg/mL) in 1X PBS 7.4, 0.1% BSA, 2 mM DTT, 2 mM $MgCl_2$ for 1 hour at room temperature. The plate was read on a TECAN Spark 20 M plate reader using the TR-FRET mode with the following parameters: Lag time: 70 μs, Integration time: 500 μs, Read A: Ex 320(25) nm (filter), Em 610(20) nm (filter), Gain 130, Read B: Ex 320(25) nm (filter), Em 665(8) nm (filter), Gain 165. FRET Signal was calculated as the ratio of [Read B]/[Read A].

#### *Steady-state GTPase assay*

WT Gαs was diluted to 6 μM (4X) in GTPase assay buffer (20 mM HEPES 7.5, 150 mM NaCl, 1 mM $MgCl_2$). The protein was 1:1 (v:v) diluted with 4X cyclic peptide stock in GTPase assay buffer, and incubated at 37 °C for an hour. The samples were then 1:1 (v:v) diluted with reaction buffer (20 mM HEPES 7.5, 150 mM NaCl, 1 mM $MgCl_2$, and 1 mM GTP) and incubated at 37 °C. After 30, 50, 70, 90 minutes, 50 μL of the sample was removed to measure the inorganic phosphate (Pi) concentration by PiColorLock™ Phosphate Detection kit. A standard curve was made using the 0.1 mM Pi stock in the kit.

#### *GDP dissociation assay*

Gα proteins were diluted to 400 nM in the EDTA buffer (20 mM HEPES 7.5, 150 mM NaCl, 1 mM EDTA-Na 8.0, 2 mM DTT). $[^3H]$GDP (1 mCi/mL, 25.2 μM) was added to a final concentration of 1.2 μM, followed by cyclic peptides addition. After incubation at 20 °C for 30 minutes, the same volume of assay buffer (20 μM HEPES-Na 7.5, 150 mM NaCl, 2 mM $MgCl_2$, and 1 mM GDP) was added to initiate $[^3H]$GDP dissociation. At various points, 10 μL of the sample was removed and mixed with 390 μL of ice-cold wash buffer (20 mM HEPES 7.5, 150 mM NaCl, 20 mM $MgCl_2$). The mixture was immediately filtered through a mixed cellulose membrane (25 mm, 0.22 μm) held by a microanalysis filter holder (EMD Millipore). The membrane was washed by ice-cold wash buffer (500 μL x 3), put in a 6-mL plastic vial and air-dried (room temperature 1.5 h). 5 mL of CytoScint-ES Liquid Scintillation Cocktail was added to each vial. After incubation overnight at room temperature, the vial was used for liquid scintillation counting with a LS 6500 Multi-Purpose Scintillation Counter. The GDP dissociation curves were fitted by the software GraphPad Prism using the following equation to calculate the dissociation rates ($k_{off}$):

$$Y=Y0 * \exp(-k_{off} * X)$$

in which ‘‘Y’’ is the radioactivity (Counts per minute) of the sample at time ‘‘X’’ (minutes), and Y0 is the calculated radioactivity of the sample at the time point 0.

#### *GTPγS binding assay*

Gα proteins were diluted to 10 μM with dilution buffer (20 mM HEPES 7.5, 150 mM NaCl, 1 mM $MgCl_2$, 2 mM DTT, and 20 μM GDP) and incubated with 5X stocks of cyclic peptides at room temperature for 2 hours. GTPγS binding was initiated by mixing with the reaction buffer at room temperature (50 nM [$^{35}$S]GTPγS and 100 μM GTPγS in dilution buffer) at room temperature. At various time points, 10 μL of the sample was removed and mixed with 390 μL of ice-cold wash buffer (20 mM HEPES 7.5, 150 mM NaCl, 20 mM $MgCl_2$). The mixture was filtered through a mixed cellulose membrane (25 mm, 0.22 μm). The membrane was washed by ice-cold wash buffer (500 μL x 3), put in a 6-mL plastic vial and air-dried (room temperature 1.5 h). 5 mL of CytoScint-ES Liquid Scintillation Cocktail (MP Biomedicals) was added to each vial. After incubation overnight at room temperature, the vial was used for liquid scintillation counting with a LS 6500 Multi-Purpose Scintillation Counter. A standard curve was generated using [35S]GTPγS. The radioactive activity (Counts per minute) of the samples were converted to the GTPγS concentration. The GTPγS binding curves were fitted by the software GraphPad Prism using the following equation to calculate the apparent GTPγS binding rates ($k_{app}$):

$$Y = Plateau * (1-exp(-k_{app} * X))$$

in which ''Y'' is the concentration of GTPγS that bound to Gα protein at time ''X'' (minutes).

#### *FRET based Gα/Gβγ interaction assay*

Biotinylated avi-Gαs (6-end, WT) and avi-Gαi (FL, WT) were diluted to 32 nM (8X) using assay buffer (1X PBS 7.4, 2 mM DTT, 0.1% BSA, 2 mM $MgCl_2$, 0.05% Tween plus 0.5 mM GDP), followed by mixing with a same volume of 8X streptavidin XL665 stock (32 nM in the assay buffer). 8X His-Gβ/γ (C68S) stock (16 nM) and 8X anti-6His-Tb cryptate stock (0.4 μg/mL) were added into the Gα/XL665 mixtures. Finally, 2X stocks of cyclic peptides were added with the protein mixtures. After incubation at room temperature for 2 hour at room temperature. The plate was read on a TECAN Spark 20 M plate reader using the TR-FRET mode with the following parameters: Lag time: 70 μs, Integration time: 500 μs, Read A: Ex 320(25) nm (filter), Em 610(20) nm (filter), Gain 130, Read B: Ex 320(25) nm (filter), Em 665(8) nm (filter), Gain 165. FRET Signal was calculated as the ratio of [Read B]/[Read A].

#### *Crystallization*

GN13/GppNHp/Gαs complex: Wild type Gαs (residues 7-380) that was preloaded with GppNHp and purified by gel filtration was concentrated to 10 mg/mL. The protein was then mixed with 1 mM of GppNHp (50 mM stock in $H_2O$) and 0.42 mM of the cyclic peptide GN13 (14 mM stock in DMSO). For crystallization, 0.2 μL of the protein sample was mixed with 0.2 μL of the well buffer containing 0.1 M HEPES 7.2, 20% PEG4000, 10% v/v 2-propanol. Crystals were grown at 20 °C in a 96-well plate using the hanging-drop vapour-diffusion method, transferred to a cryoprotectant solution (0.1 M HEPES 7.2, 20% PEG4000, 10% v/v 2-propanol, 150 mM NaCl, 20 mM HEPES 8.0, 5 mM $MgCl_2$, 1 mM GppNHp, 25% v/v glycerol), and flash-frozen in liquid nitrogen.

GD20/GDP/Gαs complex: Wild type Gαs (NCBI Reference Sequence: NP_536351.1, residues 35-380) was preloaded with GDP, purified by gel filtration and then concentrated to 11.6 mg/mL. Before crystallization, the protein was mixed with 5 mM of Dithiothreitol (0.5 M stock in $H_2O$), 1 mM of GDP (50 mM stock in $H_2O$) and 0.76 mM of the cyclic peptide GD20 (42.6 mM stock in DMSO). For crystallization, 1.5 µL of the protein sample was mixed with 1.5 µL of the well buffer containing 0.1 M Tris 8.2, 26% PEG4000, 0.8 M LiCl. Crystals were grown at 20 °C in a 15-well plate using the hanging-drop vapour-diffusion method, and flash-frozen in liquid nitrogen.

*Data collection and structure determination*

The data set was collected at the Advanced Light Source beamline 8.2.1 with X-ray at a wavelength of 0.999965 Å. Then the data set was integrated using the HKL2000 package (Otwinowski and Minor, 1997), scaled with Scala (Evans., 2006) and solved by molecular replacement using Phaser (McCoy et al., 2007) in CCP4 software suite (Winn et al., 2011). The crystal structure of GDP-bound human Gαs R201C/C237 mutant (PDB code: 6AU6) was used as the initial model. The structure was manually refined with Coot (Emsley et al., 2010) and PHENIX (Adams et al., 2010). Data collection and refinement statistics are shown in Table S1 and S2.

*Chloroalkane penetration assay (CAPA)*

The cell lines used for CAPA were HeLa cell lines, generated by Chenoweth and co-workers, that stably express HaloTag exclusively in the cytosol (Peraro et al., 2018). Cells were seeded in a 96-well plate the day before the experiment at a density of $4 \times 10^4$ cells per well. The day of the experiment the media was aspirated, and 100 µL of cyclic peptide dilutions in DMEM were added to the cells. Plate was incubated for 19.5 h at 37 °C with 5% $CO_2$. The contents of the wells were aspirated off, and wells were washed using fresh Opti-MEM for 15 min. The wash was aspirated off, and the cells were chased using 5 µM ct-TAMRA for 15 min, except for the No-ct-TAMRA control wells, which were incubated with Opti-MEM alone. The contents of the wells were aspirated and washed with fresh Opti-MEM for 30 min. After aspiration, cells were rinsed once with phosphate-buffered saline (PBS). The cells were then trypsinized, quenched with DMEM, resuspended in PBS, and analyzed using a benchtop flow cytometer (CytoFLEX, Beckman).

*BRET2 based Gα Gβγ interaction assay*

The plasmids encoding M2R was a gift from Dr. Roderick Mackinnon. The plasmids encoding Gα-RLuc8, Gβ1, and Gγ1-GFP2 were gifts from Dr. Bryan Roth. The plasmid encoding Gγ2-GFP2 was generated by replacing the Gγ1 sequence of pcDNA3.1-GGamma1-GFP2 by digestion with BamHI/XbaI and subsequent insertion of the Gγ2 sequence(MASNNTASIAQARKLVEQLKMEANIDRIKVSKAADLMAYCEAHAKEDPLLT PVPASENPFREKKFFCAIL). All plasmids were sequenced to ensure their identities.

The BRET2 assay was conducted as reported (Olsen et al., 2020). Cells were plated in 10 cm dishes at 3 million cells per dish the night before transfection. Cells were transfected using a 6:6:3:1 DNA ratio of receptor:Gα-RLuc8:Gβ:Gγ-GFP2 (750:750:375:125 ng for 10 cm dishes). Transit 2020 was used to complex the DNA at a ratio of 3 µL Transit per µg DNA, in OptiMEM (Gibco-ThermoFisher) at a concentration of 10 ng DNA per µl OptiMEM. 16 hours after transfection, cells were harvested from the plate using TrypLE and plated in poly-D-lysine-coated white, clear-bottom 96-well assay plates (Greiner Bio-One) at a density of 30,000 cells per well.

8 hours after plating in 96-well assay plates, media was replaced with 100 µL of cyclic peptide dilutions in DMEM with 1% dialyzed FBS. 16 hours after drug treatment at 37 °C with 5% $CO_2$, white backings (PerkinElmer) were applied to the plate bottoms, and growth medium was carefully aspirated and replaced immediately with 60 µL of drug dilutions in assay buffer (1× Hank's balanced salt solution (HBSS) + 20 mM HEPES, pH 7.4), followed by a 10 µl addition of freshly prepared 50 µM coelenterazine 400a. After a 5 min equilibration period, cells were treated with 30 µL of GPCR agonist or DMSO dilutions in assay buffer for an additional 5 min. Plates were then read in a TECAN Spark 20 M plate reader with 395 nm (RLuc8-coelenterazine 400a) and 510 nm (GFP2) emission filters, at integration times of 1 s per well. Plates were read serially six times, and measurements from the fourth read were used in all analyses. BRET2 ratios were computed as the ratio of the GFP2 emission to RLuc8 emission.

#### *Chemical stability assay in DMEM with 10% FBS or human Plasma*

These assays were conducted by Pharmaron Beijing CO., Ltd. Cyclic peptides working solutions were prepared at 10 µM in DMEM with 10% FBS (Avantor, Cat# 76294-180) or human plasma (Pooled, Male & Female, BioIVT, Cat# HMN666664). The assays were performed in duplicate. Vials were incubated at 37°C at 60 rpm in a water bath and taken at designated time points including 0, 480, 1080 and 1440 min. For each time point, the initiation of the reaction was staggered so all the time points were terminated with cold acetonitrile containing internal standards (IS, 100 nM alprazolam, 200 nM labetalol, 200 nM Imipramine and 2 µM ketoplofen) at the same time. Samples were vortexed then centrifuged at 4°C to remove proteins. The supernatants from centrifugation were diluted by ultra-pure $H_2O$ and used for LC-MS/MS analysis. All calculations were carried out using GraphPad Prism. Remaining percentages of parent compounds at each time point were estimated by determining the peak area ratios from extractedion chromatograms.

#### *Chemical synthesis*

##### *Solid phase synthesis of cyclic peptides*

Macrocyclic peptides (25 µmol scale) were synthesized by a standard Fmoc solid phase peptide synthesis method using a Syro Wave automated peptide synthesizer (Biotage) (Morimoto et al., 2012). After addition of a chloroacetyl group onto the N-terminal amide group (for the formation of cyclic peptide), peptides were cleaved from the NovaPEG Rink Amide resin (Novabiochem) by a solution of 92.5% trifluoroacetic acid (TFA), 2.5% 3,6-Dioxa-1,8-octanedithiol ethanedithiol (DODT), 2.5% triisopropylsilane (TIPS) and 2.5% water and precipitated by diethyl ether. To conduct the macrocyclization reaction, the peptide pellet was dissolved in 10 ml DMSO containing 10 mM tris(2-carboxyethyl)phosphine hydrochloride (TCEP), adjusted to pH>8 by addition of triethylamine (TEA) and incubated at 25 °C for 1 hour. This cyclization reaction was quenched by acidification of the solution with TFA. The crude products were purified by reverse-phase HPLC (RP-HPLC) (Shimadzu) with a Chromolith RP-18 100-25 prep column. Molecular masses were verified by a time-of-flight mass spectrometer (Waters Xevo G2-XS), and the purity was verified by analytical HPLC on a Waters Acquity UPLC BEH C18 1.7 μm column.

*General synthesis route of chloroalkane tagged cyclic peptides*

In this work, we prepared a chloroalkane tag (ct) that has been previously used with the HaloTag system (Neklesa et al., 2011). Instead of using the Rink amide resin, peptides were synthesized using the Fmoc-Wang resin (Anaspec, AS-20058) to generate a carboxylate at the C-terminus. To cap the C-terminus with the chloroalkane tag (ct), 10 equiv of chloroalkane tag (ct), 5 equiv of HATU, and 20 equiv of DIPEA were dissolved in DMF and stirred for 1 hour at room temperature. Crude peptides were purified by reverse-phase HPLC (Waters XBridge C18 column 5 μm particle size 30 x 250 mm, 5–95% acetonitrile–water + 0.1% formic acid, 40 min, 20 mL/min) to afford the chloroalkane tagged peptides.

*Characterization Data for Cyclic Peptides*

Mass Spectrometry

GN13: HRMS (ESI): Calcd for ($C_{79}H_{106}N_{16}O_{21}S$ + 2H)$^{2+}$: 824.3798, Found: 824.3973.

GD20: HRMS (ESI): Calcd for (C90H126N22$O_{20}$S + 2H)$^{2+}$: 934.4698, Found: 934.4844.

GD20-F10L: HRMS (ESI): Calcd for ($C_{87}H_{128}N_{22}O_{20}S$ + 2H)$^{2+}$: 917.4776, Found: 917.4901.

GD20-F5A: HRMS (ESI): Calcd for ($C_{84}H_{122}N_{22}O_{20}S$ + 2H)$^{2+}$: 896.4542, Found: 896.4604.

GD20-F10L/F5A: HRMS (ESI): Calcd for ($C_{81}H_{124}N_{22}O2_{0}S$+ 2H)$^{2+}$: 879.4620, Found: 879.4648.

ct-GN13-E3Q: HRMS (ESI): Calcd for ($C_{89}H_{126}ClN_{17}O_{22}S$ + 2H)$^{2+}$: 926.9416, Found: 926.9422.

ct-GD20: HRMS (ESI): Calcd for ($C_{100}H_{145}ClN_{22}O_{22}S$ + 2H)$^{2+}$: 1037.5235, Found: 1037.5303.

ct-GD20-F10L: HRMS (ESI): Calcd for ($C_{97}H_{147}ClN_{22}O_{22}S$ + 2H)$^{2+}$: 1020.5313, Found: 1020.5193.

Absorbance was recorded at 280 nm (Figure S1D-K).

### QUANTIFICATION AND STATISTICAL ANALYSIS

All of the curves in Figures except those from the BLI experiments were fitted by GraphPad Prism. Raw kinetic data collected from the BLI experiments were processed with the Data Analysis software provided by the manufacturer. All the details can be found in the figure legends and in the METHOD DETAILS. The data collection and refinement statistics of the crystal structures can be found in Table S1 and S2 (related to Figure 3 and 5, see also Figure S3 and S5).

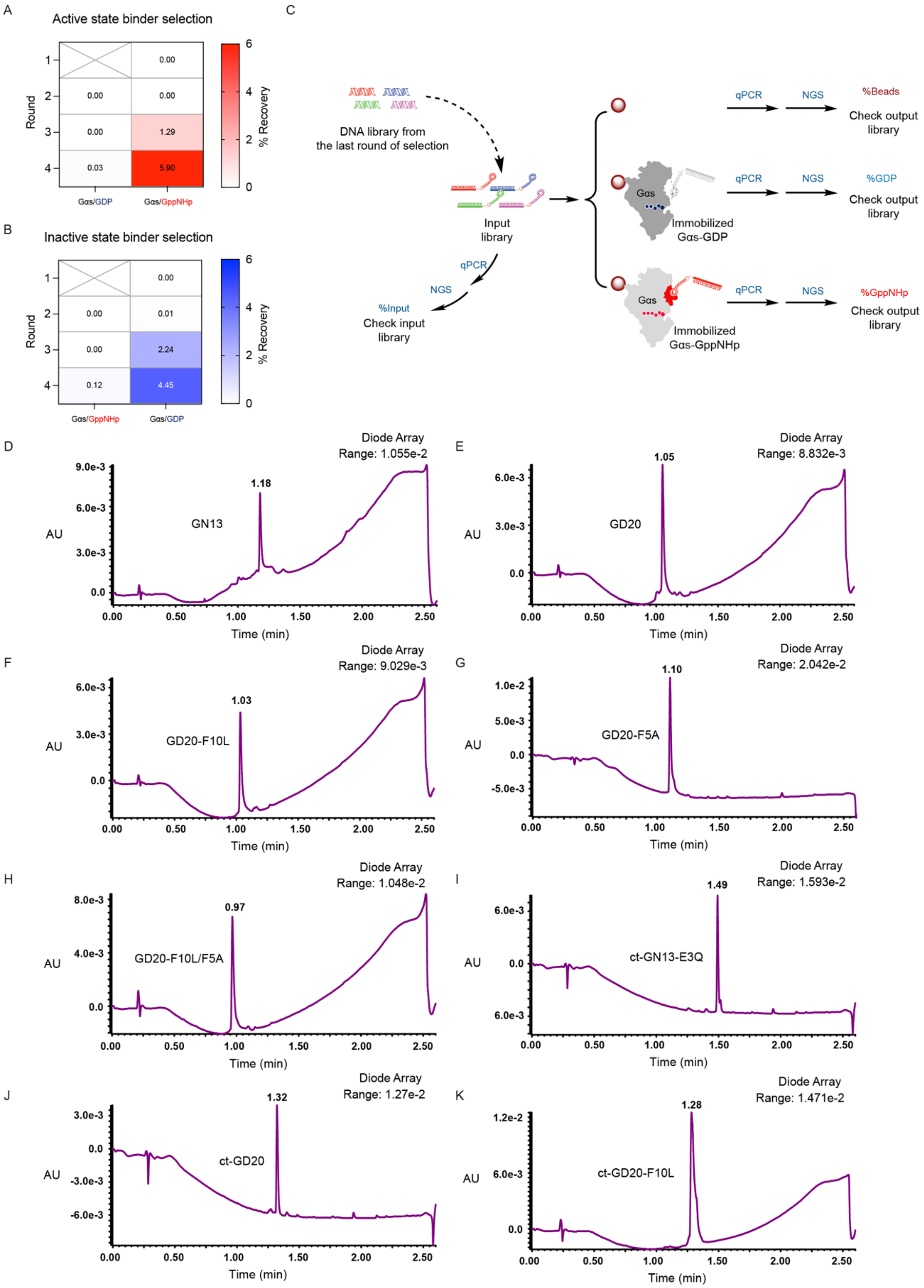

A
Active state binder selection
Round
% Recovery
Gαs/GDP
Gαs/GppNHp
B
Inactive state binder selection
C
DNA library from the last round of selection
Input library
qPCR
NGS
%Input
Check input library
%Beads
Check output library
Immobilized Gαs-GDP
%GDP
Immobilized Gαs-GppNHp
%GppNHp
D
GN13
E
GD20
F
GD20-F10L
G
GD20-F5A
H
GD20-F10L/F5A
I
ct-GN13-E3Q
J
ct-GD20
K
ct-GD20-F10L
Diode Array
Time (min)
AU

### Figure S1. RaPID selection of state-selective Gαs binding cyclic peptides, related to Figure 1

(A and B) The percentage of enriched peptide-mRNA-cDNA complex in the input library after each selection was quantified by qPCR. Cyclic peptides that bind to GppNHp-bound (A) or GDP-bound (B) Gαs were enriched through four rounds of RaPID selection. To ensure a maximum library diversity at the initial stage of selection, negative selection was not included in the first round of selection. (C) Comparison selection. DNA sequences of cyclic peptide binders from the last round of selection were quantified and identified by qPCR and next generation sequencing (NGS). A peptide-mRNA-cDNA complex library was produced based on the above-mentioned DNA sequences and equally split into three fractions. Binding of each individual peptide-mRNA-cDNA complex to blank, GDP-bound Gαs-immobilized or GppNHp-bound Gαs-immobilized beads was quantified by qPCR and NGS, respectively. (D-K) Analytical HPLC Traces of resynthesized cyclic peptides. Absorbance was recorded at 280 nm. GN13 (D). GD20 (E). GD20-F10L (F). GD20-F5A (G). GD20-F10L/F5A (H). GN13-E3Q (I). ct-GD20 (J). ct-GD20-F10L (K).

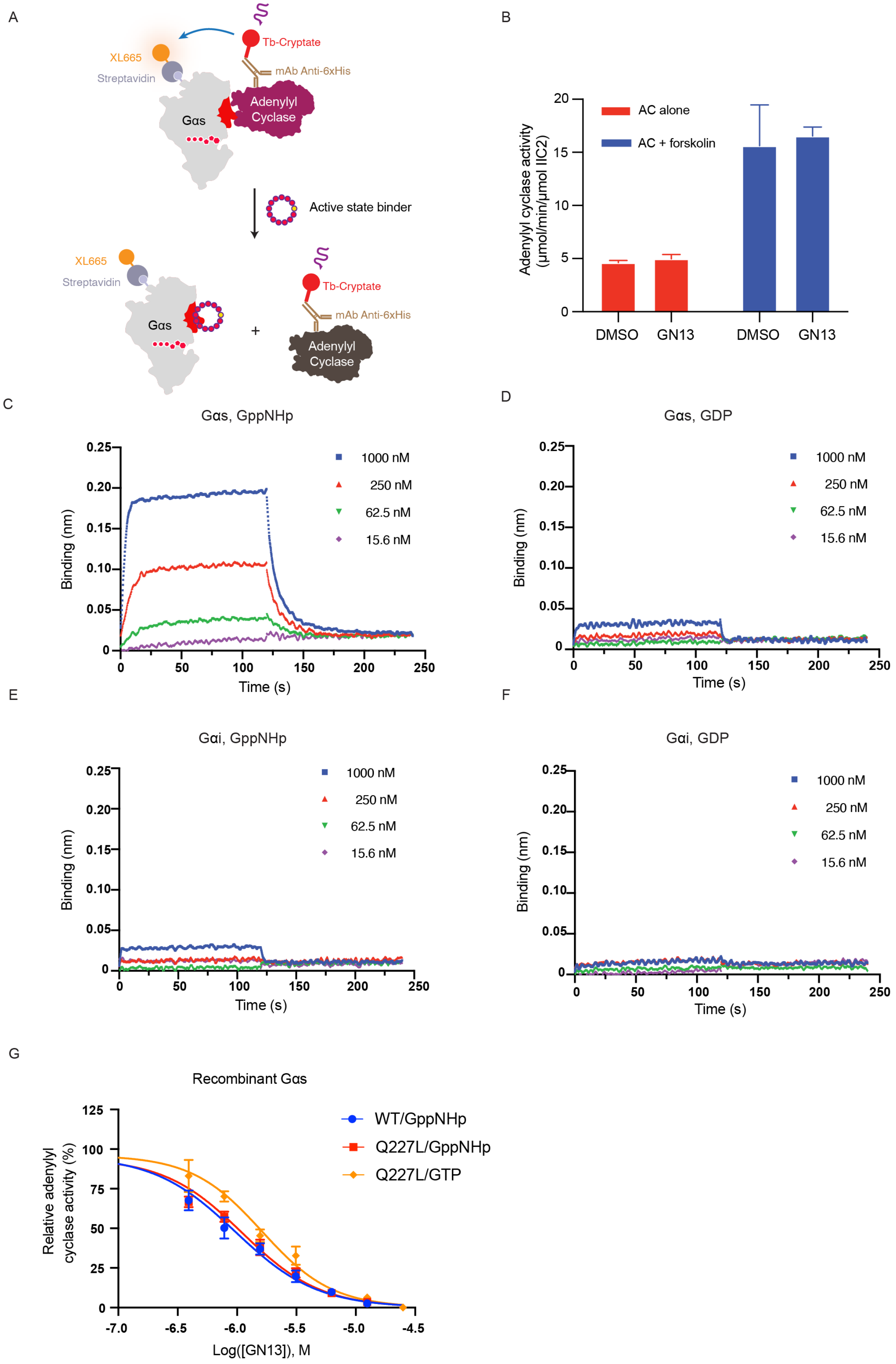
A
Tb-Cryptate
XL665
Streptavidin
mAb Anti-6xHis
Adenylyl Cyclase
Gαs
Active state binder
XL665
Streptavidin
Tb-Cryptate
mAb Anti-6xHis
Gαs
+
Adenylyl Cyclase
B
AC alone
AC + forskolin
Adenylyl cyclase activity (μmol/min/μmol IIC2)
20
15
10
5
0
DMSO
GN13
DMSO
GN13
C
Gαs, GppNHp
Binding (nm)
0.25
0.20
0.15
0.10
0.05
0
1000 nM
250 nM
62.5 nM
15.6 nM
0
50
100
150
200
250
Time (s)
D
Gαs, GDP
Binding (nm)
1000 nM
250 nM
62.5 nM
15.6 nM
Time (s)
E
Gαi, GppNHp
Binding (nm)
1000 nM
250 nM
62.5 nM
15.6 nM
Time (s)
F
Gαi, GDP
Binding (nm)
1000 nM
250 nM
62.5 nM
15.6 nM
Time (s)
G
Recombinant Gαs
Relative adenylyl cyclase activity (%)
125
100
75
50
25
0
WT/GppNHp
Q227L/GppNHp
Q227L/GTP
-7.0
-6.5
-6.0
-5.5
-5.0
-4.5
Log([GN13]), M

### Figure S2. Biochemical characterization of GN13, related to Figure 2

(A) Schematic representation of active state binders inhibiting the protein-protein interaction between biotinylated Gαs WT and His-tagged adenylyl cyclase.
(B) GN13 did not directly inhibit adenylyl cyclase activity in the absence of Gαs. 25 μM of GN13 or DMSO were mixed with adenylyl cyclase (VC1/IIC2), followed by addition of DMSO or forskolin. After adding ATP, the reaction was carried out at 30 °C for 10 min. Production of cAMP was evaluated by the LANCE Ultra cAMP kit. The data represent the mean ± SE of three independent measurements.
(C to F) Binding kinetics of GN13 to Gα proteins were quantified using bio-layer Interferometry. The assay was performed in duplicate, and the data represent one of the two replicates. Biotinylated Gα proteins were immobilized to give a relative intensity of 3nm on streptavidin biosensors. Association (t = 0-120 s) and dissociation (t = 120-240 s) cycles of compounds were started by dipping sensors into GN13 dilutions and control buffer. (C) GN13 binding to GppNHp-bound Gαs. (D) GN13 binding to GDP-bound Gαs. (E) GN13 binding to GppNHp-bound Gαi. (F) GN13 binding to GDP-bound Gαi.
(G) Activation of adenylyl cyclase by Gαs oncogenic mutants was inhibited by GN13 in a dose-dependent manner.

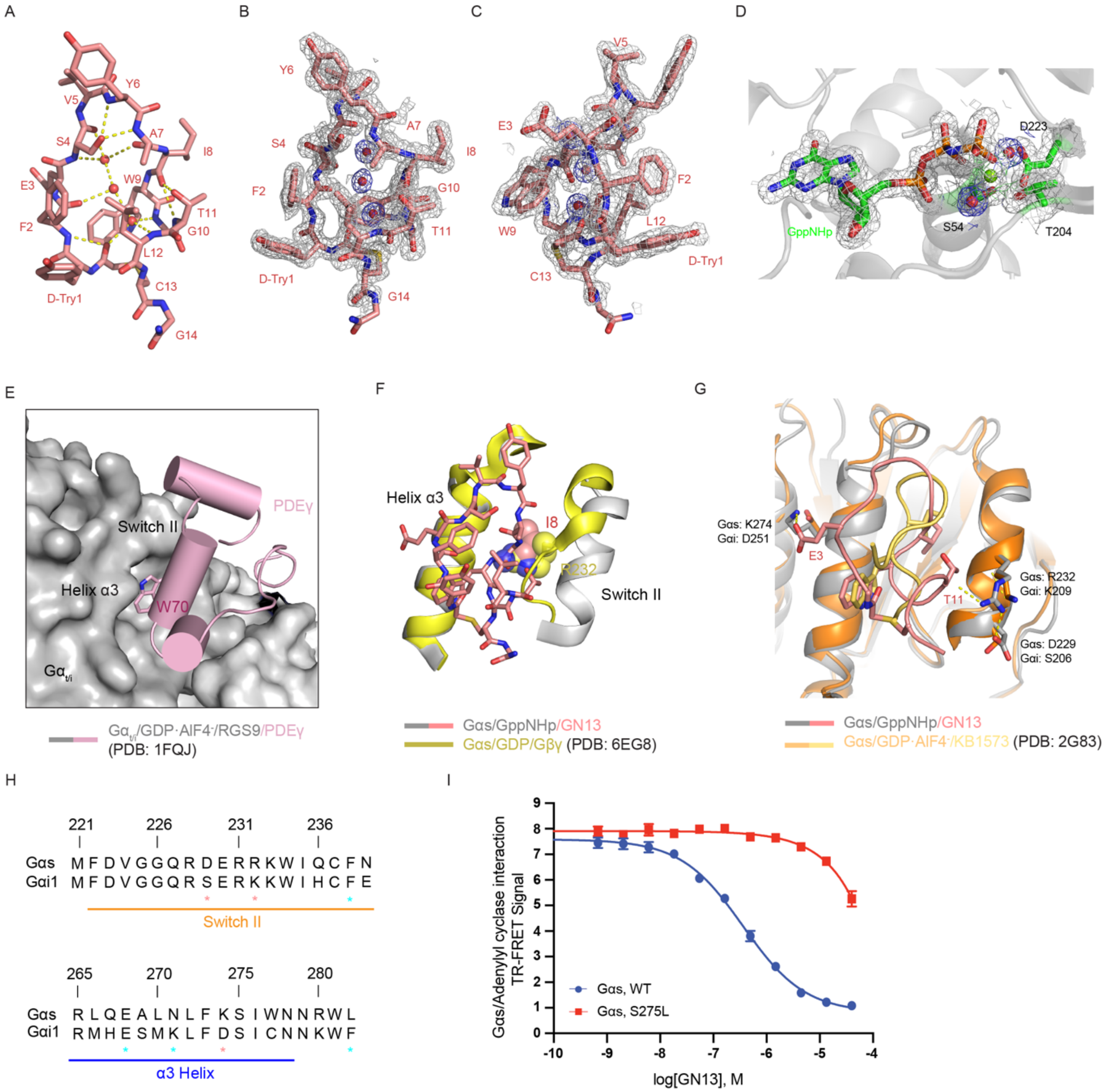

A
B
C
D
E
F
G
H
I
GppNHp
D223
S54
T204
PDEγ
Switch II
Helix α3
W70
Helix α3
I8
R232
Switch II
Gαs: K274
Gαi: D251
E3
T11
Gαs: R232
Gαi: K209
Gαs: D229
Gαi: S206
Gαt/i/GDP·AlF4-/RGS9/PDEγ (PDB: 1FQJ)
Gαs/GppNHp/GN13
Gαs/GDP/Gβγ (PDB: 6EG8)
Gαs/GppNHp/GN13
Gαs/GDP·AlF4-/KB1573 (PDB: 2G83)
221 226 231 236
Gαs M F D V G G Q R D E R R K W I Q C F N
Gαi1 M F D V G G Q R S E R K K W I H C F E
Switch II
265 270 275 280
Gαs R L Q E A L N L F K S I W N N R W L
Gαi1 R M H E S M K L F D S I C N N K W F
α3 Helix
Gαs/Adenylyl cyclase interaction TR-FRET Signal
log[GN13], M
Gαs, WT
Gαs, S275L

**Figure S3. GN13 specifically inhibits Gαs through binding to a crystallographically defined pocket, related to Figure 3**

(A) GN13 adopts a highly ordered three-dimensional structure through intramolecular and intermolecular hydrogen bonding network. GN13 is shown as salmon sticks. Three water molecules with well-defined electron density are shown as red spheres. Hydrogen bonds are represented by yellow dash lines.
(B and C) Electron density map of GN13. GN13 is shown as salmon sticks. Three water molecules with well-defined electron density are shown as red spheres. The 2mFo-DFc electron density map of the structure is contoured at 1.0 σ and colored grey (GN13) and blue (Water), respectively.
(D) Electron density map of GppNHp. GppNHp and the side chains of S54, T204 and D223 are shown as sticks. The $Mg^{2+}$ and two water molecules coordinated with the $Mg^{2+}$ are shown as green and red spheres, respectively. The 2mFo-DFc electron density map of the structure is contoured at 1.0 σ.
(E) Structure of the GDP•$AlF4^-$-bound $Gα_{t/i}$/RGS9/PDEγ complex (PDB: 1FQJ). A critical tryptophan residue from PDEγ (pink, cartoon) engages the hydrophobic pocket between the switch II region and the α3 helix. $Gα_{t/I}$ and RGS9 are shown as surface. PDEγ is shown as cartoon.
(F) Structural basis for nucleotide-state-selective binding of GN13 to Gαs. In GDP-bound Gαs, switch II is partially disordered, which disrupts polar contacts with GN13 and creates extensive steric hindrance. In particular, R232 of switch II (shown in space filling) is predicted to create a steric clash with I8 of GN13.
(G) Structural basis for G protein class-specific binding of GN13 to Gαs. Gαs interacts with GN13 though two specific charge interactions. However, Gαi misses those critical GN13-binding residues.
(H) Sequence alignment of Gα proteins around the cyclic peptide binding site. The residue numbering is based on Gαs. α3 helix and Switch II regions are indicated with colored lines. The residues that appear to determine the specificity of GN13 (salmon) or GD20 (cyan) are shown with asterisks.
(I) GN13 inhibited the protein-protein interaction of Gαs WT with adenylyl cyclase in a dose-dependent manner (blue). This inhibitory effect was significantly diminished by the S275L mutation (red). The data represent the mean ± SD of three independent measurements.

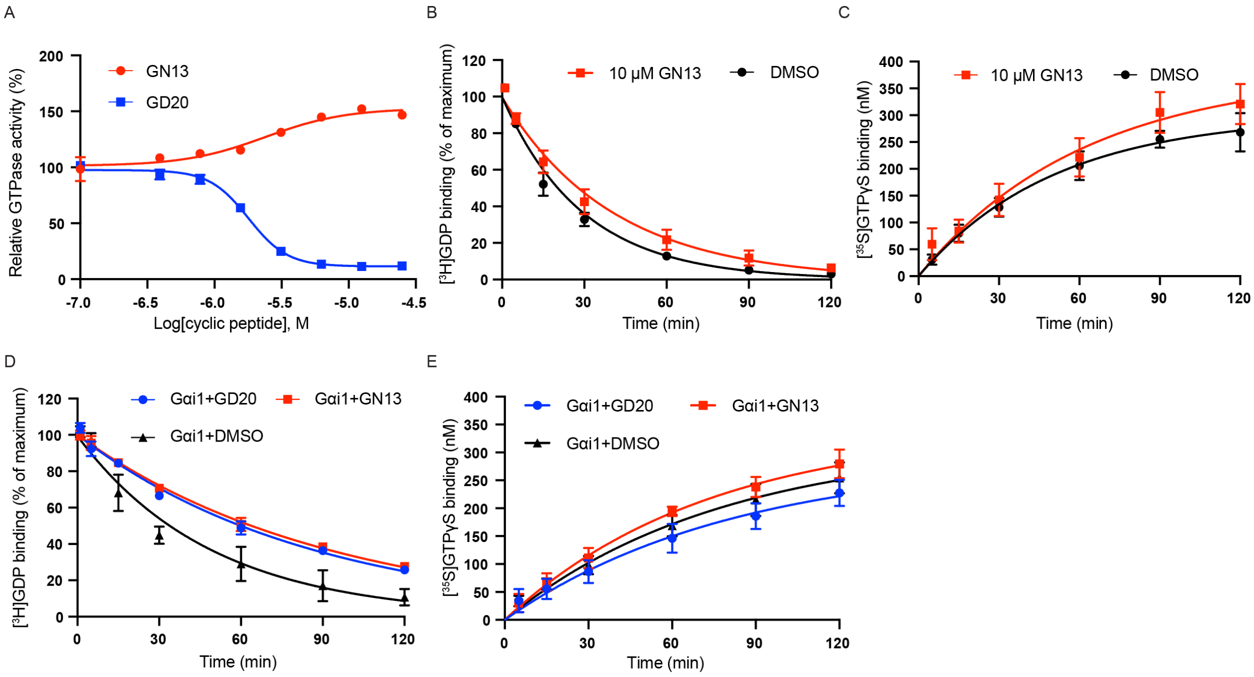
A
GN13
GD20
Relative GTPase activity (%)
Log[cyclic peptide], M
B
10 μM GN13
DMSO
[3H]GDP binding (% of maximum)
Time (min)
C
10 μM GN13
DMSO
[35S]GTPγS binding (nM)
Time (min)
D
Gαi1+GD20
Gαi1+GN13
Gαi1+DMSO
[3H]GDP binding (% of maximum)
Time (min)
E
Gαi1+GD20
Gαi1+GN13
Gαi1+DMSO
[35S]GTPγS binding (nM)
Time (min)

**Figure S4. GN13 and GD20 modulate Gαs GTPase activity through a Gαs-specific manner, related to Figure 4**

(A) Gαs steady-state GTPase activity was modulated by GN13 and GD20 in a dose-dependent manner. The data represent the mean ± SD of three independent measurements.
(B) The rates of GDP dissociation from Gαs in the presence (red) or absence (black) of 10 μM GN13 were determined. Gαs preloaded with [$^3$H]GDP was assayed in a buffer containing 1 mM $MgCl_2$, 0.5 mM GDP, and the indicated concentration of GN13. The data represent the mean ± SD of three independent replicates.
(C) The rates of GTPγS binding to Gαs in the presence (red) or absence (black) of 10 μM GN13 were determined by mixing GDP-bound Gαs with a mixture of [$^{35}$S] GTPγS and GTPγS in a buffer containing 1 mM $MgCl_2$. The data represent the mean ± SD of three independent replicates.
(D) The rates of GDP dissociation from Gαi1 in the presence of 10 μM GN13 (red), or 10 μM GD20 (blue) or DMSO (black) were determined. Gαi1 preloaded with [$^3$H]GDP was assayed in a buffer containing 1 mM $MgCl_2$, 0.5 mM GDP, and the indicated concentration of cyclic peptides. The data represent the mean ± SD of three independent replicates.
(E) The rates of GTPγS binding to Gαi1 in the presence of 10 μM GN13 (red), or 10 μM GD20 (blue) or DMSO (black) were determined by mixing GDP-bound Gαi1 with a mixture of [$^{35}$S] GTPγS and GTPγS in a buffer containing 1 mM $MgCl_2$. The data represent the mean ± SD of three independent replicates.

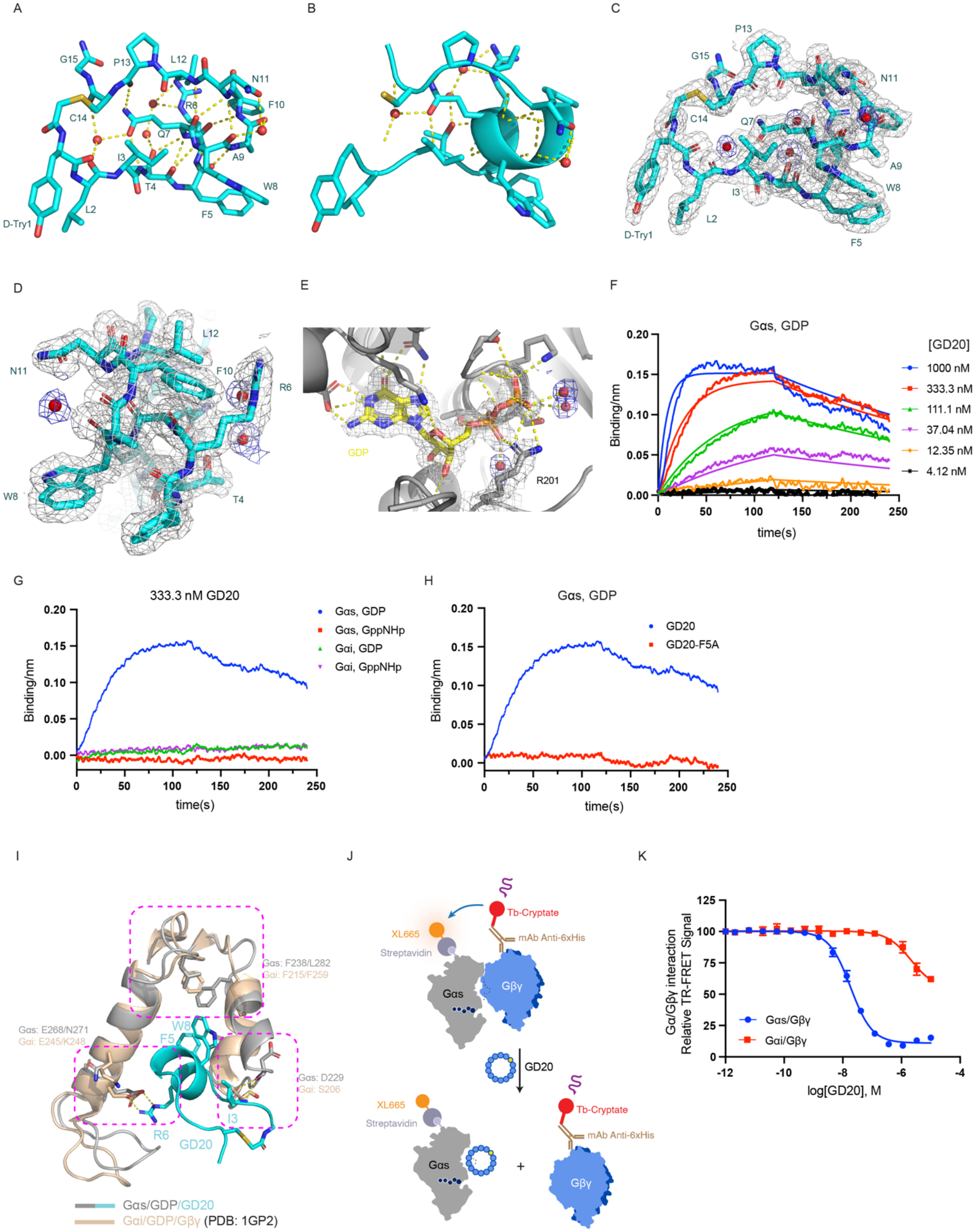

A
G15
P13
L12
N11
R6
F10
C14
Q7
I3
A9
T4
W8
F5
L2
D-Try1
B
C
P13
G15
N11
C14
Q7
A9
I3
W8
L2
D-Try1
F5
D
L12
N11
F10
R6
W8
T4
E
GDP
R201
F
Gαs, GDP
[GD20]
1000 nM
333.3 nM
111.1 nM
37.04 nM
12.35 nM
4.12 nM
Binding/nm
time(s)
G
333.3 nM GD20
Gαs, GDP
Gαs, GppNHp
Gαi, GDP
Gαi, GppNHp
Binding/nm
time(s)
H
Gαs, GDP
GD20
GD20-F5A
Binding/nm
time(s)
I
Gαs: F238/L282
Gαi: F215/F259
Gαs: E268/N271
Gαi: E245/K248
W8
F5
Gαs: D229
Gαi: S206
I3
R6
GD20
Gαs/GDP/GD20
Gαi/GDP/Gβγ (PDB: 1GP2)
J
Tb-Cryptate
XL665
mAb Anti-6xHis
Streptavidin
Gαs
Gβγ
GD20
XL665
Tb-Cryptate
Streptavidin
mAb Anti-6xHis
Gαs
+
Gβγ
K
Gα/Gβγ interaction
Relative TR-FRET Signal
Gαs/Gβγ
Gαi/Gβγ
log[GD20], M

### Figure S5. GD20 specifically inhibits Gαs through binding to a crystallographically defined pocket, related to Figure 5

(A and B) GD20 adopts a highly ordered three-dimensional structure through intramolecular and intermolecular hydrogen bonding network. GD20 is shown as cyan sticks (A) or cartoon (B). Four water molecules with well-defined electron density are shown as red spheres. Hydrogen bonds are represented by yellow dash lines.
(C and D) Electron density map of GD20. GD20 is shown as cyan sticks. Four water molecules with well-defined electron density are shown as red spheres. The 2mFo-DFc electron density map of the structure is contoured at 1.0 σ and colored grey (GD20) and blue (Water), respectively.
(E) Electron density map of GDP. GDP and the side chain of R201 are shown as sticks. The $Mg^{2+}$ and two water molecules coordinated with the $Mg^{2+}$ are shown as green and red spheres, respectively. The 2mFo-DFc electron density map of the structure is contoured at 1.0 σ.
(F to H) Binding kinetics of GD20 and GD20-F5A to Gα proteins were quantified using bio-layer Interferometry. The assay was performed in duplicate, and the data represent one of the two replicates. Biotinylated Gα proteins were immobilized to give a relative intensity of 2.5nm on streptavidin biosensors. Association (t = 0-120 s) and dissociation (t = 120-240 s) cycles of compounds were started by dipping sensors into cyclic peptide dilutions and control buffer. (F) GD20 binding to GDP-bound Gαs. (G) 333.3 nM of GD20 binding to different Gα proteins. (H) 333.3 nM of GD20 or GD20-F5A binding to GDP-bound Gαs.
(I) Structural basis for G protein class-specific binding of GD20 to Gαs. Gαs interacts with GD20 though three major specificity-determining sites. However, Gαi misses those critical GD20-binding residues.
(J) Schematic representation of inactive state binders inhibiting the protein-protein interaction between biotinylated Gαs WT and His-tagged Gβγ(C68S).
(K) GD20 inhibited the protein-protein interaction between biotinylated Gαs WT and His-tagged Gβγ(C68S) in a dose-dependent manner. GD20 was 100-fold more selective for Gαs than Gαi. The data represent the mean ± SD of three independent replicates.

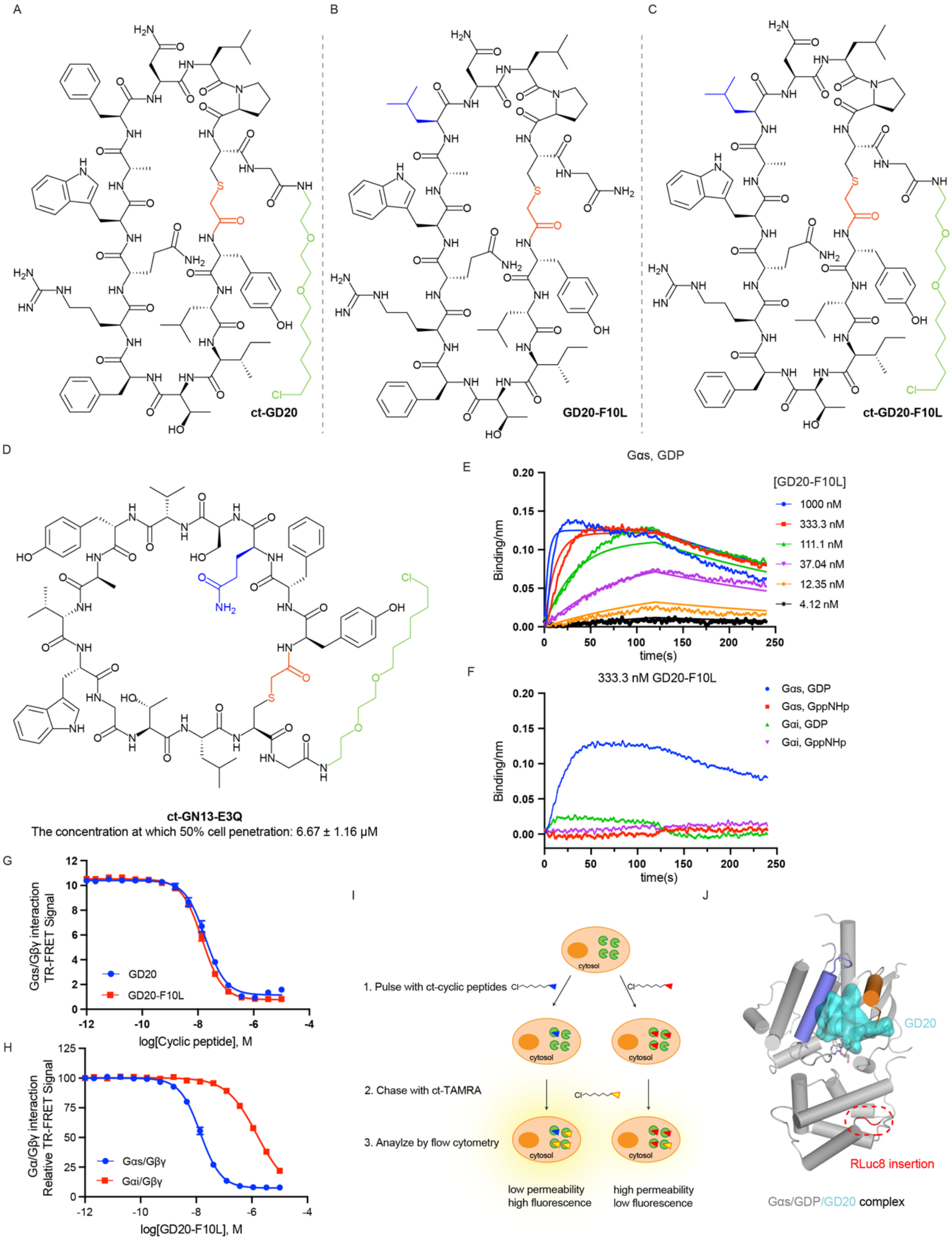
A
ct-GD20
B
GD20-F10L
C
ct-GD20-F10L
D
ct-GN13-E3Q
The concentration at which 50% cell penetration: 6.67 ± 1.16 μM
E
Gαs, GDP
[GD20-F10L]
1000 nM
333.3 nM
111.1 nM
37.04 nM
12.35 nM
4.12 nM
Binding/nm
time(s)
F
333.3 nM GD20-F10L
Gαs, GDP
Gαs, GppNHp
Gαi, GDP
Gαi, GppNHp
Binding/nm
time(s)
G
Gαs/Gβγ interaction TR-FRET Signal
GD20
GD20-F10L
log[Cyclic peptide], M
H
Gα/Gβγ interaction Relative TR-FRET Signal
Gαs/Gβγ
Gαi/Gβγ
log[GD20-F10L], M
I
1. Pulse with ct-cyclic peptides
2. Chase with ct-TAMRA
3. Anaylze by flow cytometry
cytosol
low permeability
high fluorescence
high permeability
low fluorescence
J
GD20
RLuc8 insertion
Gαs/GDP/GD20 complex

**Figure S6. A cell permeable GD20 analog F10L specifically inhibits Gαs/Gβγ interaction through a Gαs-specific manner, related to Figure6**

(A to D) Structure of derivatized cyclic peptides. (A) ct-GD20 (B) GD20-F10L (C) ct-GD20-F10L (D) ct-GN13-E3Q. Mutations are colored blue. The ct tag is colored green. Cell penetration of ct-GN13-E3Q was also measured using the CAPA assay.
(E to F) Binding kinetics of GD20-F10L to different Gα proteins were quantified using bio-layer Interferometry. The assay was performed in duplicate, and the data represent one of the two replicates. Biotinylated Gα proteins were immobilized to give a relative intensity of 2.5nm on streptavidin biosensors. Association (t = 0-120 s) and dissociation (t = 120-240 s) cycles of compounds were started by dipping sensors into cyclic peptide dilutions and control buffer. (E) GD20-F10L binding to GDP-bound Gαs. (F) 333.3 nM of GD20-F10L binding to different Gα proteins.
(G to H) GD20-F10L inhibited the protein-protein interaction between biotinylated Gαs WT and His-tagged Gβγ(C68S) in a dose-dependent manner (G). GD20-F10L was 100-fold more selective for Gαs than Gαi (H). The data represent the mean ± SD of three independent replicates.
(I) Schematic representation of the chloroalkane penetration assay (CAPA). HeLa cells stably express GFP-tagged HaloTag on the mitochondrial outer membrane. If the pre-dosed chloroalkane-tagged molecule (ct-molecule) penetrates the cell membrane, it will covalently label HaloTag and block subsequent HaloTag labeling with ct-TAMRA. Intracellular ct-TAMRA fluorescence intensity is inversely related to the amount of cytosolic ct-molecule.
(J) The GD20/Gαs complex structure provides structural basis for the Rluc8 insertion. Rluc8 is inserted between αB and αC helices.

**Table S1: Data collection and refinement statistics for the Gαs/GppNHp/GN13 complex. Related to Figure 3**

| | Gαs/GppNHp/GN13 complex |
|---|---|
| **Data collection** | |
| Space group | P 21 21 21 |
| Cell dimensions | |
| *a*, *b*, *c* (Å) | 68.905, 78.332, 80.043 |
| *α*, *β*, *γ* (°) | 90, 90, 90 |
| Resolution (Å) | 50.00-1.57 (1.60-1.57)[a] |
| $R_{merge}$, $R_{meas}$, and $R_{pim}$ | 0.074 (1.040), 0.086 (0.972), 0.023 (0.405) |
| $I/\sigma(I)$ | 27.3 (1.32) |
| $CC_{1/2}$ | 0.997 (0.783) |
| Completeness (%) | 96.9 (71.2) |
| Total reflections | 58703 |
| Unique reflections | 57627 |
| Redundancy | 12.4 (4.0) |
| **Refinement** | |
| Resolution (Å) | 43.45-1.574 |
| No. reflections | 51161 |
| $R_{work}$ | 0.1970 |
| $R_{free}$ | 0.2291 |
| No. atoms | |
| Protein | 3079 |
| Ligand/ion (specify/describe) | 44 |
| Water | 132 |
| *B* factors | |
| Protein | 21.98 |
| Ligand/ion | 13.41 |
| Water | 22.43 |
| R.m.s. deviations | |
| Bond lengths (Å) | 0.014 |
| Bond angles (°) | 1.39 |
| Ramachandran analysis | |
| Favored (%) | 98.91 |
| Allowed (%) | 0.82 |
| Outliers (%) | 0.27 |
| Rotamer outliers (%) | 0.60 |
| Clashscore | 3.57 |

[a] Values in parentheses are for highest-resolution shell.

**Table S2: Data collection and refinement statistics for the Gαs/GDP/GD20 complex. Related to Figure 5**

| | Gαs/GDP/GD20 complex |
|---|---|
| **Data collection** | |
| Space group | P1 |
| Cell dimensions | |
| *a*, *b*, *c* (Å) | 58.105, 81.771, 76.912 |
| *α*, *β*, *γ* (°) | 81.266, 83.844, 90.698 |
| Resolution (Å) | 50.00-1.95 (1.98-1.95)[a] |
| $R_{merge}$, $R_{meas}$, and $R_{pim}$ | 0.081 (1.202), 0.113 (1.260), 0.059 (0.762) |
| $I/\sigma(I)$ | 11.40 (0.815) |
| $CC_{1/2}$ | 0.991 (0.422) |
| Completeness (%) | 97.0 (87.7) |
| Total reflections | 99153 |
| Unique reflections | 95974 |
| Redundancy | 3.4 (2.0) |
| **Refinement** | |
| Resolution (Å) | 48.54-1.95 |
| No. reflections | 82224 |
| $R_{work}$ | 0.2178 |
| $R_{free}$ | 0.2580 |
| No. atoms | |
| Protein | 11279 |
| Ligand/ion | 132 |
| Water | 329 |
| *B* factors | |
| Protein | 29.36 |
| Ligand/ion | 18.22 |
| Water | 21.27 |
| R.m.s. deviations | |
| Bond lengths (Å) | 0.004 |
| Bond angles (°) | 0.70 |
| Ramachandran analysis | |
| Favored (%) | 97.45 |
| Allowed (%) | 2.25 |
| Outliers (%) | 0.30 |
| Rotamer outliers (%) | 0.98 |
| Clashscore | 3.86 |

[a] Values in parentheses are for highest-resolution shell.

**Table S3: Kinetics analysis of cyclic peptides-Gαs interaction by BLI. Related to Figure 2D, 4C, S2, S5 and S6**

| | $K_D$ (nM) | $K_{on}$ ($M^{-1}s^{-1}$) | $K_{off}$ ($s^{-1}$) |
|---|---|---|---|
| GN13/GppNHp/Gαs | 190 ± 16 | $2.71 \times 10^5 \pm 2.13 \times 10^4$ | $5.13 \times 10^{-2} \pm 1.31 \times 10^{-3}$ |
| GD20/GDP/Gαs | 31.4 ± 0.7 | $1.09 \times 10^5 \pm 1.28 \times 10^3$ | $3.43 \times 10^{-3} \pm 6.97 \times 10^{-5}$ |
| GD20-F10L/GDP/Gαs | 14.5 ± 0.4 | $2.47 \times 10^5 \pm 3.71 \times 10^3$ | $3.58 \times 10^{-3} \pm 8.50 \times 10^{-5}$ |

**Table S4: Chemical stability of Gαs binding cyclic peptides in DMEM with 10% FBS. Related to STAR Methods**

| | Half-life (hour) [a] |
|---|---|
| GN13 | >76 |
| GD20 | >142 |
| GD20-F10L | 37.37 - 50.32 |

[a] Values represent 95% confidence intervals
The data were analyzed from two independent replicates.

**Table S5: Plasma stability of Gαs binding cyclic peptides. Related to STAR Methods**

| | Half-life (hour) [a] |
|---|---|
| GN13 | >82 |
| GD20 | 16.37 - 27.13 |
| GD20-F10L | 7.16 - 12.74 |

[a] Values represent 95% confidence intervals
The data were analyzed from two independent replicates.
